# Neuroanatomy of stomatopod central complexes offers putative neural substrate for oriented behaviors in crustaceans

**DOI:** 10.1101/2022.06.10.495695

**Authors:** Alice Chou, Marcel E. Sayre, Chan Lin, Thomas W. Cronin

**Affiliations:** Department of Biological Sciences, University of Maryland Baltimore County, Baltimore, Maryland 21250; Department of Neuroscience, University of Arizona, Tucson, AZ 85721; Department of Invertebrate Zoology, Smithsonian National Museum of Natural History, Washington, D.C. 20560

**Keywords:** stomatopod, crustacean, central complex, lateral complex, neuroanatomy

## Abstract

All insects studied to date possess a centrally located group of neuropils, known collectively as the central complex, that has been implicated in sensory integration and motor action selection. Among the functions prescribed to the central complex, none is perhaps as intriguing as its role in orientation and navigation. Neurobiological correlates of both current and desired headings have been described in insect CXs. Despite the diversity of arthropods, understanding of the CX as a navigational center originates entirely from terrestrial insects. Stomatopod crustaceans, commonly referred to as mantis shrimps, form an order of predatory marine crustaceans with intricate and diverse visual systems that maintain the distinction of being the only fully aquatic animal known to utilize the navigational strategy of path integration. They utilize idiothetic, celestial, and landmark cues to orient in the benthos. Here, we investigate the neuroanatomy of adult and developing mantis shrimp central complexes and associated neuropils to begin understanding this brain region in a sensorially and behaviorally complex crustacean.

## Introduction

The central complex (CX) is a prominent group of modular neuropils found across the midline of all insect brains studied to date. It is proposed to underlie the neural basis of insect behavior, in which the selection of actions is driven by the integration of sensory information, prior experiences, and internal state. The CX is strongly implicated in fundamental and higher-order functions such as sensory integration, flight and walking coordination, courtship, sleep, visual and spatial memory, and attention (Liu et al, 2006; Martin et al., 2015; Ofstad et al., 2011; Pfeiffer and Homberg, 2014; Turner-Evans and Jayaraman, 2016; Donlea et al., 2014; Guo and Ritzmann, 2013; Ilius et al., 2007; Popov et al., 2003; Sakai and Kitamoto, 2006; Strauss, 2002; Weir et al., 2014; Turner-Evans et al., 2017). Perhaps the best studied function of the central complex is its role in insect orientation and navigation. In many insects, allocentric celestial cues, such as the sky polarization pattern and positions of the sun or moon, are relayed to the CX via the anterior visual pathway where they are ultimately integrated to form an internal representation of heading direction (Homberg et al., 2011; Beetz and el Jundi, 2018; Dacke et al., 2011; el Jundi et al., 2019; el Jundi et al. 2015; Heinze and Homberg, 2007; Sauman et al., 2005; Honkanen et al., 2019; Pegel et al., 2019; Turner-Evans et al., 2020). Additionally, a separate but parallel network of cells in the CX have recently been shown to encode an internal representation of traveling direction in flies (Lu et al., 2022; Lyu et al., 2022). This network has further been proposed to provide the neural basis for path integration, or vector-based navigation, and has been successfully modeled from the bee CX (Stone et al., 2017; see also Hulse et al. 2021 for modeling in the fly).

Anatomically, the insect CX is located in the protocerebrum, the most anterior neuromere of the fused cerebral ganglion. In all pterygote insects described to date, the CX is comprised of the protocerebral bridge (PB), the fan-shaped body (FB) and the ellipsoid body (EB) which together comprise the central body (CB), as well as the paired globular noduli (NO) (Pfeiffer and Homberg, 2014; Strausfeld, 2012). These neuropils are characteristically organized into discrete internal layers and columns, which have been respectively suggested to represent spatial directions and cue information (Pfeiffer and Homberg, 2014). The CX is connected to many parts of the surrounding protocerebrum, most prominently to the lateral complex (LX), which in insects is composed of the paired lateral accessory lobes (LALs), the gall (GA), the bulbs (BUs), and several other areas (Hulse et al., 2021). However, beyond the handful of insects in which the CX has been described in detail, comparatively little is known about other arthropod CXs, including those of crustaceans.

Malacostracan crustaceans make up an exceptionally large and diverse group of arthropods found in marine, estuarine, freshwater, and terrestrial habitats ranging from the Arctic Ocean to hydrothermal vents. Deep examinations of crustacean neurobiology have generated important models for our understanding of nervous system function and organization. For example, studying the lobster stomatogastric nervous system has provided insights into the principles of neuromodulation; and electrical synapses were first described as part of the crayfish escape circuit (Harris-Warrick and Marder, 1991; Marder, 2012; Furschpan and Potter, 1959). Despite the undeniable contributions of crustaceans to neuroscience and their potential for future discoveries, the central complex is relatively unexplored in these arthropods.

While the brains of insects and crustaceans share common developmental origins, crustacean CX structures are typically more simplified (Stollewerk and Simpson, 2005; Strausfeld, 2012). In decapod crustaceans, for example the crayfish *Cherax descructor*, the CX is composed of a small, bilateral PB that sits above a thin, bi-layered CB. The upper and lower divisions of the CB are suggested to be comparable to the insects FB and EB, respectively (Homberg, 2008; Utting et al., 2000).

Stomatopod crustaceans, also known as mantis shrimp, form one of the most charismatic orders within Malacostraca and have attracted significant attention for their intricate and diverse visual systems and ballistic strikes. They are benthic predators, found around the globe, that diverged from their closest crustacean relatives about 340 million years ago (Van Der Wal et al., 2017). There are over 400 known species within the six extant superfamilies (Squilloidea, Gonodactyloidea, Bathysquilloidea, Lysiosquilloidea, Hemisquilloidea, and Parasquilloidae) that occupy environments from brightly lit intertidal zones to the deep sea (Porter et al., 2010).

Stomatopods have an uncanny ability to manipulate their environment (for food acquisition, burrow construction, as well as inter- and intraspecific interactions) and possess specialized eyes that are capable of color vision, linear and circular polarization vision, ultraviolet vision, and motion vision (Chiou et al., 2008; Cronin and Marshall, 2001; Cronin and Marshall, 1989a; Cronin and Marshall, 1989b; Cronin et al., 2014; Marshall, 1988; Marshall and Land, 1993; Marshall et al., 1991a; Marshall et al., 1991b). Recent behavioral evidence in *Neogonodactylus oerstedii* has also shown stomatopods to be adept navigators and the only fully aquatic animals thus far known to path integrate (Patel and Cronin, 2020a). Small burrows and cavities provide safety and shelter for most stomatopods. However, they must leave these homes to forage for food and find mates. Because this leaves them vulnerable to predation or aggression from conspecifics, stomatopods must be able to quickly return home when circumstance demands. They do so using landmarks for place identification, celestial cues for knowledge of their current orientation, and path integration informed by a ranked hierarchy of cues (Patel and Cronin, 2020a,b,c). While detailed navigational experiments in other crustaceans are lacking, these navigational strategies strikingly resemble those used by terrestrial insects, including desert ants, and dung beetles (Müller and Wehner, 1988; Rossel and Wehner, 1984; Dacke et al, 2003; Byrne et al. 2003).

In accordance with their sensory and behavior abilities, stomatopods possess elaborate neural architecture. Of the few studies so far conducted on stomatopod neurobiology, most have centered upon understanding how their extensive visual channels are processed in the optic lobes (Kleinlogel and Marshall, 2006; Kleinlogel and Marshall, 2005; Kleinlogel et al., 2003; Lin et al., 2019; Lin and Cronin, 2018; Thoen et al, 2017a). However, recent work has found that stomatopod CX neural architecture may be more similar to that of dicondylic insects, than of other malacostracan crustaceans (Thoen et al., 2017b). The present study further characterizes the stomatopod CX and associated neuropils, including a previously undescribed ellipsoid body-like neuropil, and discusses implications of stomatopod CX neuroarchitecture in the context of development, behavior, and ecology.

## Materials and Methods

### Animals

Adult specimens of the stomatopod species *Squilla empusa* and *Neogonodactylus oerstedii* were collected by Gulf Specimen Marine Laboratories and K. B. Marine Life in the Florida Keys, USA and were shipped directly to the University of Maryland Baltimore County. Adult *Haptosquilla trispinosa*, *Chorisquilla hystrix, Pullosquilla sp.*, and stomatopod post-larvae were collected from areas near the Lizard Island Research Station, Australia (Great Barrier Reef Marine Park Permit no. G12/35005.1, Fisheries Act no. 140763). Live *H. trispinosa* were transported in fresh seawater that was refreshed daily to the University of Maryland Baltimore County. *C. hystrix* brain tissue was dissected, fixed, and stored in 0.1M phosphate-buffered saline (PBS) prior to transit.

*N. oerstedii* larvae were hatched from adults collected by K. B. Marine Life in Florida, USA. The egg masses remained with their mothers and were monitored daily. Once eyespots, a sign of imminent hatching, were visible through the eggs, cheesecloth was wrapped around the housing container to prevent any loss of larvae through water flow channels. Once the larvae hatched, they were separated into six containers with fresh artificial seawater. 50% water changes were performed each day. Larval stages were tracked by the number of days post-hatching (Manning and Provenzano, 1978; Provenzano and Manning, 1978). At each stage, starting with Stage III, several individuals were fixed in 4% paraformaldehyde with 10-12% sucrose in 0.1M PBS buffer at 4°C overnight. Samples were then washed 3 x 20 min in PBS and stored in PBS until further processing.

*Pullosquilla sp.* larvae were hatched from eggs collected from shallow coral rubble in areas near the Lizard Island Research Station, Australia. Adults were collected in mated pairs, both with and without egg masses, and kept in running natural seawater. Eggs were monitored for the development of retinal eyespots. Once hatched, larvae used in this work were removed from parental containers every 12 hours over 5 days and fixed in 4% paraformaldehyde with 10-12% sucrose in 0.1M PBS at 4°C overnight. Samples were then washed 3 x 20 min in PBS and stored in PBS until further processing.

Post-larval collections were performed at night while wading. Pelagic stomatopod larvae and post-larvae exhibit positive phototaxis. Thus, using a strong underwater light source, we attracted larvae and post-larvae at ∼1 meter depth and collected them with hand nets. Animals were then sorted into superfamilies by gross morphology and housed in natural running seawater. Every 12 hours, four to five animals from each collection trip were fixed in 4% paraformaldehyde with 10-12% sucrose at 4°C for 12-14 hours. Samples were then washed 3 x 20 min and stored in PBS until further processing.

### Immunohistochemistry

Adult *N. oerstedii* were cold-anesthetized and decapitated. Heads were partially dissected to expose nervous tissue, and then were fixed in 4% paraformaldehyde in 0.1M phosphate-buffered saline (PBS; Fisher Scientific, Fair Lawn, NJ, USA) at pH 7.4 with 10-12% sucrose. After 3 x 20 min washes in 0.1 M PBS, brain tissue was carefully dissected from the cuticle. Dissected adult brains and whole larval and post-larval samples were embedded in album gelatin blocks. Blocks were fixed overnight in 4% paraformaldehyde (Electron Microscopy Sciences, Hatfield, PA, USA) in 0.1 M PBS at 4°C before being washed with 0.1 M PBS 3 x 10 min, then sections at 60 μm or 120 μm with a vibratome. Sections were washed with PBS-TX (0.5% Triton X-100 in PBS) 6 x 20 min then blocked in 5% normal goat serum (NGS; Vector Laboratories, Burlingame, CA, USA) for 1 h before being incubated on a shaker overnight with primary antibodies (Table 1) in PBS-TX with 5% NGS. Sections were then washed in 0.1 M PBS with 5% NGS 6 x 20 min. Secondary antibody solutions were prepared by centrifuging 1000 μL of 0.1 M PBS with 5% NG with secondary IgG antibodies, including Alexa Fluor 488-, 555-, 633-, 647-, Cy3, and Cy5 goat anti-mouse and goat anti-rabbit (Invitrogen; 2.5:1000) for 13,000 rpm for 4 minutes at 4°C. Sections were incubated with the top 900 μL of the secondary antibody seolutions at room temperature on a shaker overnight. Finally, sections were rinsed in 0.1 M PBS 6 x 20 min before being mounted and coverslipped in a medium of 25% polyvinyl alcohol, 25% glycerol, and 50% PBS.

**Table 1.** List of antibodies used in immunohistochemistry experiments.

| Antibody | Supplier | Host | Dilution |
| --- | --- | --- | --- |
| alpha-tubulin | DHSB, #12G10 | Mouse (monoclonal) | 1 to 333 |
| DCO | Generous gift from Dr. Daniel Kalderon | Rabbit (polyclonal) | 1 to 200 |
| FMRFamide | Immunostar, Hudson, WI 20080 | Rabbit (polyclonal) | 1 to 200 |
| GABA | Genetex , Irvine, CA 92606 | Rabbit (polyclonal) | 1 to 200 |
| GAD | SigmaAlrich, St. Louis, MO 68178 | Rabbit (polyclonal) | 1 to 200 |
| Neuropeptide Y | Abcam, Waltham, MA 02453 | Rabbit (polyclonal) | 1 to 200 |
| RII | Generous gift from Dr. Daniel Kalderon | Rabbit (polyclonal) | 1 to 200 |
| Serotonin (5-HT) | Immunostar, Hudson, WI 20080 | Rabbit (polyclonal) | 1 to 200 |
| Synapsin<br>"SYNORF-1" | DSHB, #3C11 | Mouse (monoclonal) | 1 to 333 |

### Western blots

Western blot analysis was used to confirm the expression of DCO in *N. oerstedii* brains by comparing it to brain tissue from crayfish *Procambarus clarkii*, the fruit fly *Drosophila melanogaster,* and the cricket *Gryllodes sigillatus*. Anti-DCO was used at a concentration of 1:500.

### Bodian reduced-silver staining

Animals were cold-anesthetized and decapitated for silver staining following Bodian’s original method as described below. Brain tissue was dissected out in room temperature AAF fixative (17 ml 100% ethanol, 1 ml glacial acetic acid, 2 ml 37% formaldehyde) and fixed for 3 hours at room temperature. Tissue was washed 2 x 10 min in 70% ethanol, then dehydrated through an ethanol series (70% ethanol, 90% ethanol, 2 x 100% ethanol; 8 minutes each), and cleared in terpineol overnight at room temperature in the dark. Tissue was then transferred to a 1:1 mixture of terpineol and xylene for 5 minutes before being held in 100% xylene. Tissue was then incubated for 15 minutes in a warm mixture of 1:1 xylene and melted Paraplast Plus (Tyco, Mansfield, MA), transferred into dishes with melted paraffin to embed, and cooled quickly in an ice bath.

Blocks were trimmed, then serially sectioned at 12 mm. Sections were arranged and flattened on slides gently warmed to 50°C. Slides were incubated in xylene 2 x 10 min, rehydrated (2 x 100% ethanol, 90% ethanol, 70% ethanol, 50% ethanol, ddH_2_O; 8 minutes each), and silver impregnated overnight at 60°C in 250 mL ddH_2_O, 6 g of clean copper fillings, and 2.5 g in-house synthesized protargol (Pan et al., 2013). Tissue was developed for 5 minutes in a solution of 1% hydroquinone and 2% sodium sulfite, washed in a gentle stream of tap water for 3 minutes, then rinsed 2 x 30 seconds in ddH_2_O. Tissue was toned in a filtered 1% gold chloride solution under a strong light for 10 minutes, then washed in ddH_2_O 2 x 30 seconds. Tissue was differentiated in 2% oxalic acid for 8 minutes and fixed in 5% sodium thiosulfate. Finally, tissue was dehydrated through an ethanol series (70% ethanol, 90% ethanol, 2 x 100% ethanol; 8 minutes each) and mounted in Entellan (Electron Microscopy Science, Hatfield, PA, USA).

### Image acquisition and processing

Immunolabeled sections and whole mounts of adult, post-larval, and larval brains were imaged using a Leica SP5 laser scanning confocal microscope (Leica Microsystems, Buffalo Grove, IL, USA) or a Zeiss LSM 900 Confocal with Airyscan 2. On the former, image stacks were collected with a 10x/0.4 Plan Apochromat objective or a 20x/0.75 PL APO CS2 objective. On the Zeiss confocal microscope, image stacks were collected with either a 10x/0.45 NA DIC objective or a 20x/0.80 NA DIC objective. Both were collected at 1024 x 1024 pixel resolution at approximately 1 mm depth intervals.

Sections of *N. oerstedii* larvae were imaged with an LSM 5 Pascal confocal microscope (Zeiss, Oberkochen, Germany). Image stacks were collected with 20x/0.5 plan Neofluar or 40x/1.0 oil plan apochromat objectives at 1024 x 1024 pixel resolution at approximately 1 mm depth intervals. Selected images from all datasets were adjusted for brightness and contrast using the ImageJ implementation FIJI (general public license, downloadable from http://fiji.sc).

Light microscopy images of serially sectioned Bodian-stained brains and were either collected using a Nikon digital camera D5100 with a T-mount NDPL-1 microscope camera adapter (AmScope, Irvine, CA, USA) connected to an Olympus BH-2 microscope (Olympus, Tokyo, Japan) or with a 5 MP Axiocam 105 color mounted on a Zeiss Axio Observer 7 Inverted Microscope with Apotome 0.2 (Carl Zeiss AG, Oberkochen, Germany).

### Terminology

The terminology used in this document is consistent with the systematic nomenclature developed by the Insect Brain Name Working Group (Ito et al., 2014). Unless otherwise noted, directional terminology was determined by the longitudinal axis of the body because mantis shrimp brains do not undergo the 90° rotation seen in many hexapods. Identification of nerves and connectives follows Sandeman et al., (1992), which offers a common nomenclature for decapod crustaceans, as the guide developed for insect brains is insufficient for that purpose.

## Results

### Overview of the stomatopod central complex

The stomatopod central complex spans the sagittal midline and is bilaterally symmetric (Fig. 1 and 2). Beginning at the anterior end of the neuraxis is a conspicuous protocerebral bridge (PB) (Supp. Fig. 1). In adult stomatopods, it sits within the peripheral cell bodies on the rostral end of the cerebral ganglion (Fig. 2B-E). The neuropil extends continuously across the midline with each side curving ventrally and has the unusual addition of a pair of ‘arms’ that extend rostrally before bending back towards the dorsal side of the PB. This neuroarchitecture is consistent across at least three species (Supp. Fig. 1A-C). The arms do not appear to be FMRF-aminergic. However, antisera raised against FMRF-amide does label the midline-spanning region of the PB and a significant portion of the superior medial protocerebrum (SMP) that extend to the medial-ventral cusp of the PB (Supp. Fig. 1E-G).

**Figure 1.**
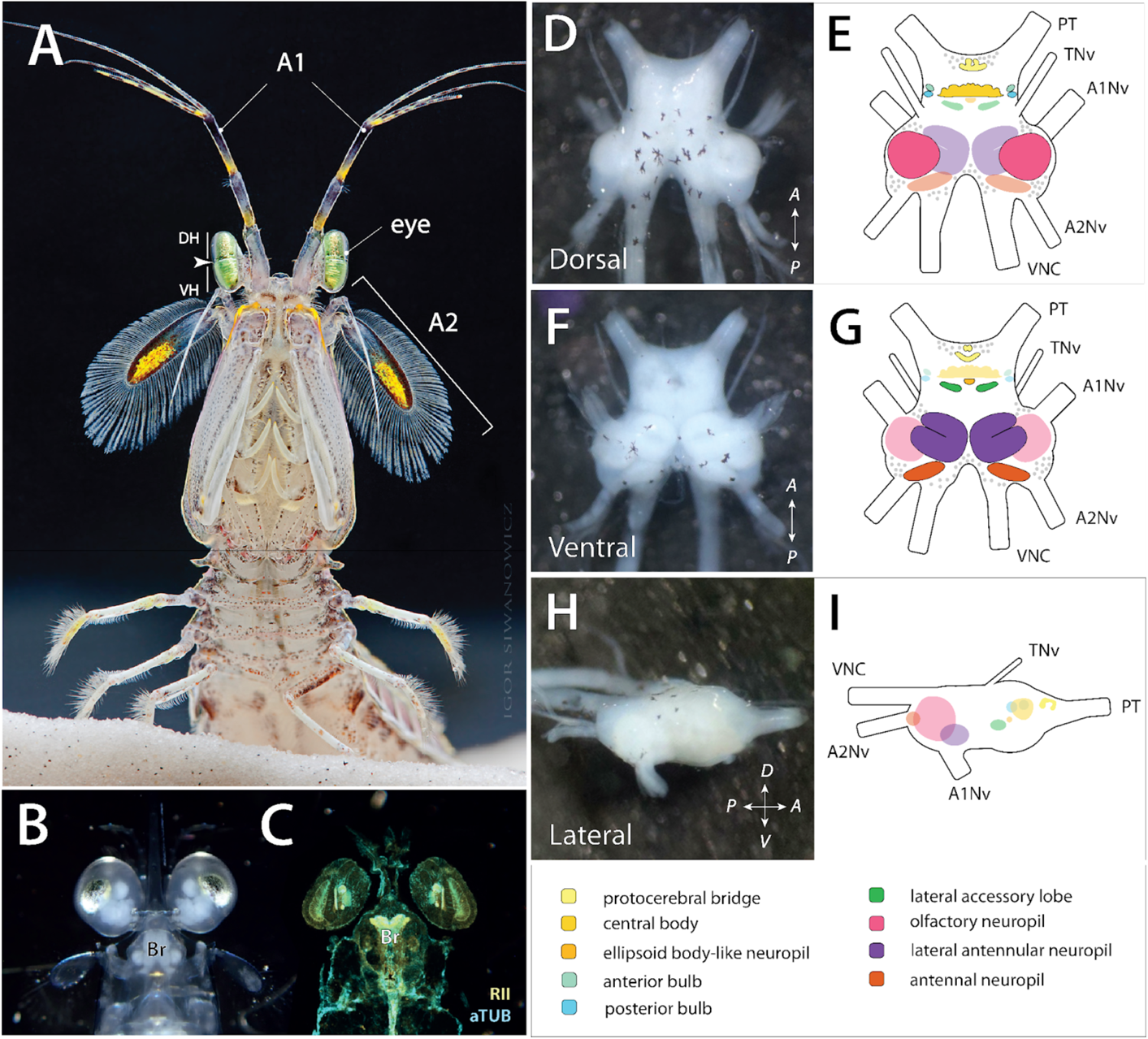
Organization and orientation of the stomatopod central brain. (**A**) Mantis shrimp possess three primary pairs of sensory appendages: two pairs of antennae, the antennules (A1) and antennae (A2), and their tripartite eyes. Photo of *Squilla empusa* by Igor Siwanowicz. (**B**) Nervous tissues, such as the optic lobes and brain (Br), are visible as white structures through the eyestalks and rostral plates of developing stomatopods, as in this ventral view of a *Lysiosquillina maculata* postlarva by Judy Jing-Wen Wang. (**C**) Double immunolabeling with anti-alpha tubulin (cyan) and anti-RII (yellow) in a *N. oerstedii* larva also clearly show the locations of neural structures. (**D**-**I**) Orientation of selected neuropils in the stomatopod brain, including central complex and lateral complex neuropils. Information travels to and from the central brain via the following connectives: the protocerebral tract (PT), which connects to the eyestalk neuropils; the tegumentary nerves (TN), that contain mechanosensory afferents from the dorsal carapace; antennujlar (antenna I) nerve (A1Nv), which carries information from the antennules; antennal (antenna II) nerve (A2Nv) from the antennae; and the ventral nerve cord (VNC). Photos by Connor Brainard.

**Figure 2.**
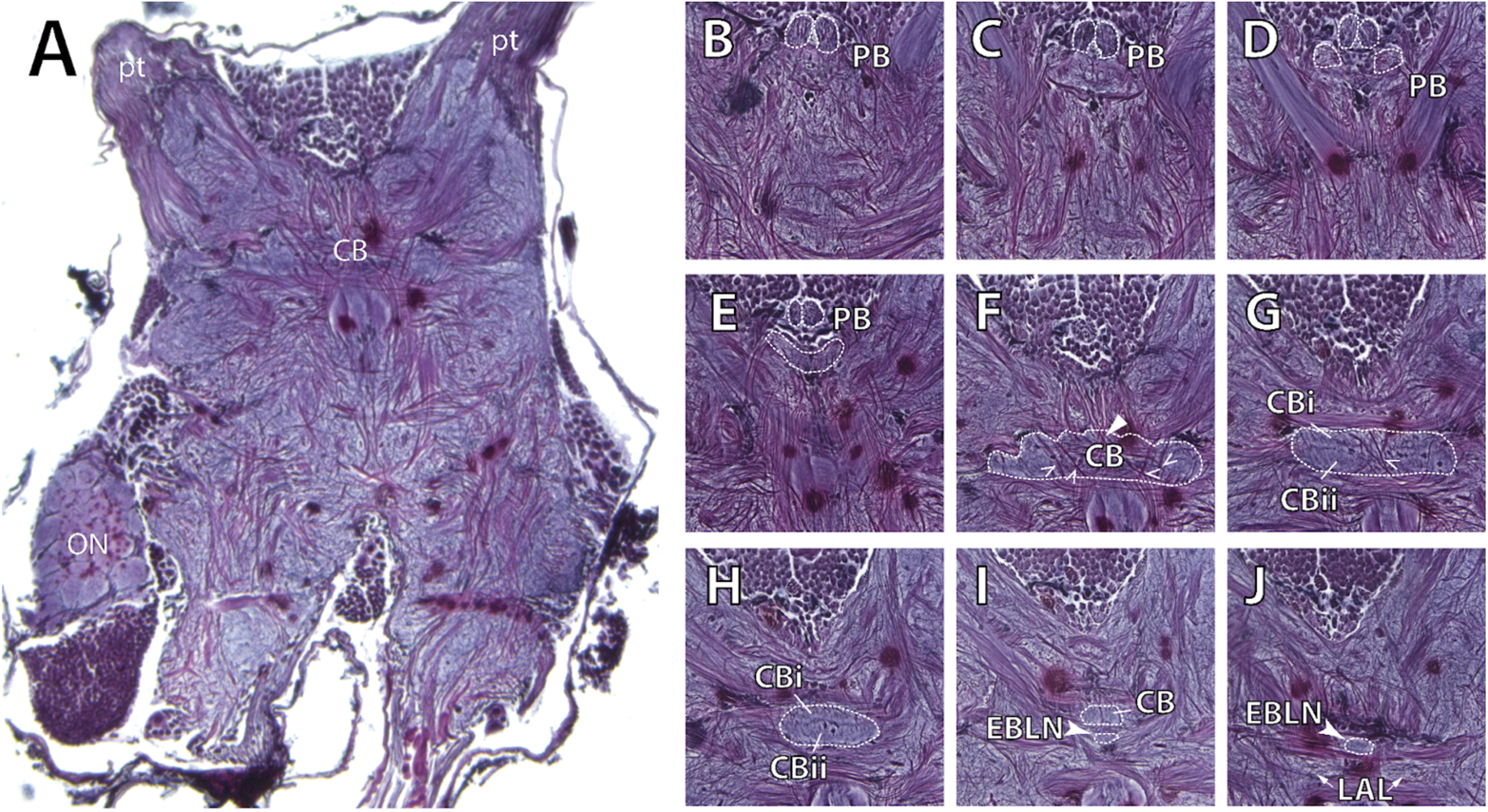
Relative anatomical positions of central complex neuropils in *N. oerstedii*. (**A**) Bodian reduced silver staining shows the central body (CB) neuropil located prominently in the protocerebrum spanning the midline. Using this method, axons are stained dark pink, while neuropils remain violet. (**B**-**J**) Dorsal to ventral serial sections of the stomatopod central complex (outlined). The protocerebral bridge (PB) consists of a midline spanning component and two arms that extend out dorsally then curve antero-ventrally. Tracts descending from the PB to the central body (CB) from the anterior chiasma (carets) and the posterior chiasma (triangle). The CB is divided into an upper (CBi) and lower (CBii) division. Immediately anterior and ventral to the CB is the ellipsoid body-like neuropil (arrowhead; EBLN).

As described in Thoen et al. (2017b), four pairs of axon bundles called the *w*-, *x*-, *y*-, and *z*-tracts connect the PB to a large, bi-layered central body (CB). The fibers that give rise to these bundles extend from CX columnar neurons towards the main body of the PB. These bundles completely decussate, in which the axon bundles from each hemisphere cross over each into the other. This is in contrast to other crustaceans in which only a portion of the axons supplied from the PB to the CB cross over (Thoen et al., 2017b; Strausfeld, 2012). As in insects, these neurons give rise to the posterior chiasma between the PB and CB and the anterior chiasma between the CB and lateral complex (LX).

The stomatopod CB is caudal to the PB and is divided into distinct upper (CBi) and lower (CBii) partitions, as resolved with anti-synapsin immunolabeling (Fig. 3). Extensive lamination of both divisions are revealed by anti-5HT immunostaining (Fig. 3C). These are possibly formed by tangential neurons, which in insects have ramifications throughout the rest of the protocerebrum (Pfeiffer and Homberg, 2014). There is notable differential immunoreactivity between CBi and CBii. Specifically, anti-synapsin labeling resolves two layers distinct within CBi, and as well as punctate staining in CBii (Fig. 3A, B). Immunolabeling with antisera raised against FMRF-amide also resolves distinct basket-like components within the CBi (Fig. 3D).

**Figure 3.**
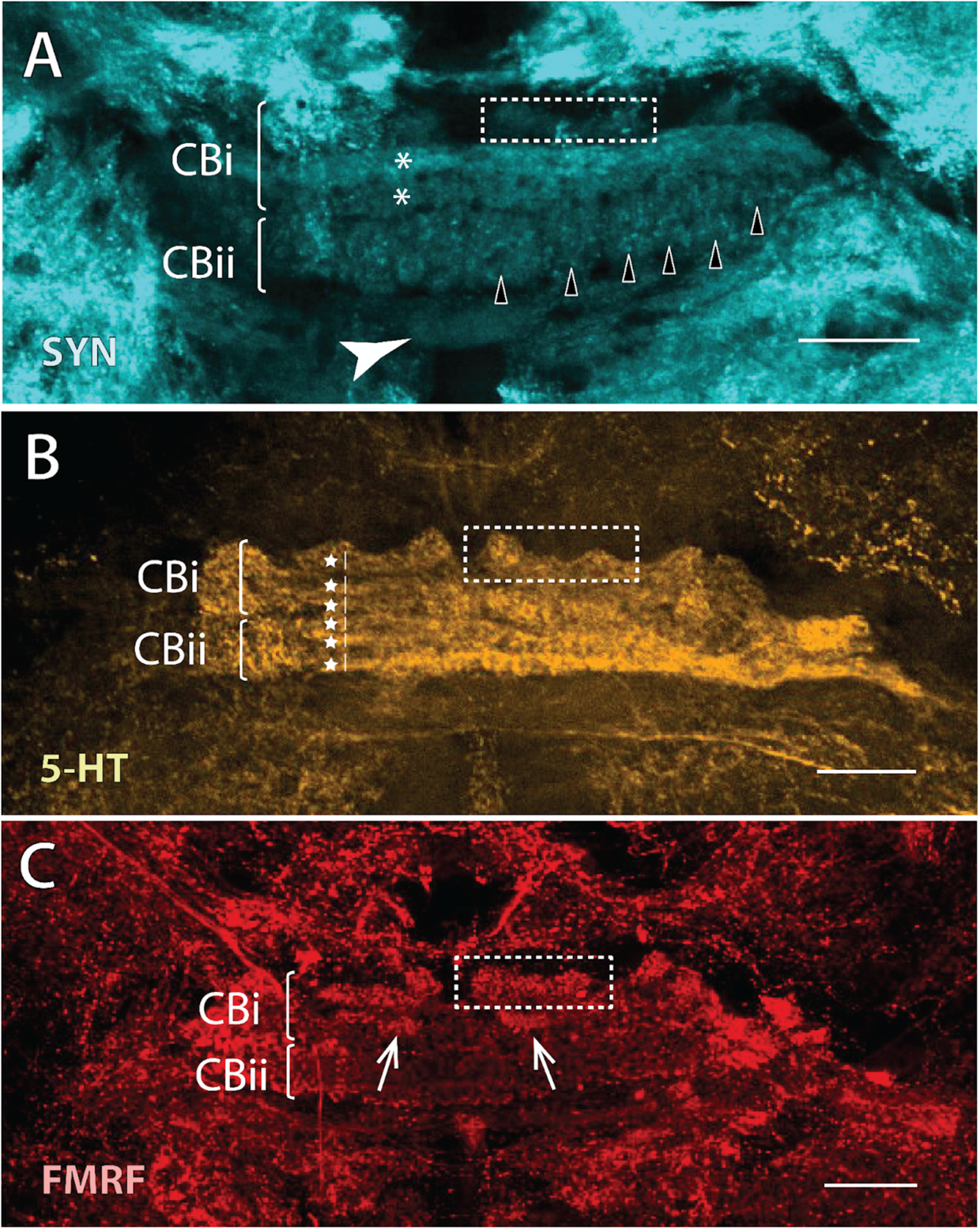
Differential immunoreactivity within the stomatopod central body. (**A-C**) The stomatopod central body is divided into two layers, the anterior (CBi) and posterior (CBii) divisions. (**A**) Anti-synapsin (SYN) reactivity shows that the CBi has at least two layers (asterisks). Additionally, the CBii has at least six columnar units on each side (right units labeled with black triangles). (**B**) Anti-serotonin (5HT) immunostaining reveals at least six distinct stratifications (stars) in the CB between both the CBi and CBii. Furthermore, defined bilateral basket-like components (right units boxed) can be identified along the rostral length of the CBi that are immunoreactive to anti-SYN (**A**), anti-5HT (**B**), and anti-FMRFamide (**C**). Ovoid structures (arrows; **C**) that are FMRF-amide positive are flank the underside of the baskets. Scale bars = 50 um.

Prior work has suggested that stomatopods are the only malacostracan crustaceans described thus far to possess the paired NO, which are a defining characteristic of pterygote insect CXs (Thoen et al., 2017b). Thoen’s study places the noduli caudo-ventral to the lower partition of the CB. However, no such structures were identified in our study.

### An ellipsoid body-like neuropil

DCO is a catalytic subunit of protein kinase A (PKA), which plays a critical role in learning and memory (Skoulakis et al., 1993). Certain regions of arthropod brains preferentially react to antisera raised against DCO, namely the mushroom bodies and the ellipsoid body (EB) (Wolff et al., 2012). Immunostaining with anti-DCO and anti-synapsin in *N. oerstedii* and *Squilla empusa*, representing different stomatopod superfamilies (Gonodactylidea and Squilloidea, respectively), reveals a bulbous neuropil that exhibits phenotypic similarity to the EB in the cockroach, *P. americana* (Fig. 4A-C). This neuropil is also strongly immunoreactive to antisera raised against RII, a regulatory subunit of PKA (Crittenden et al., 1998; Fig. 4D).

**Figure 4.**
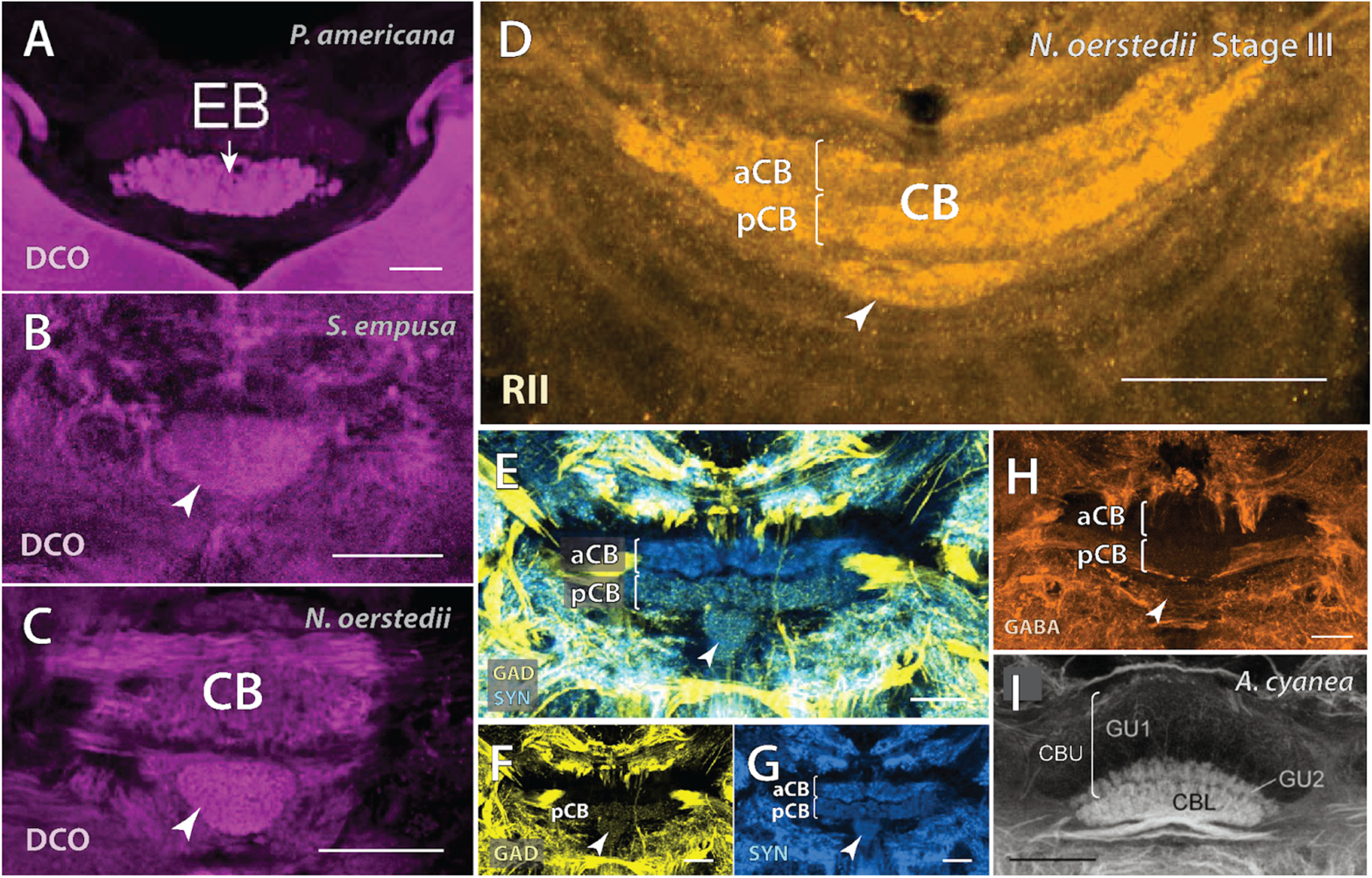
The ellipsoid body in cockroach (**A**; modified from Wolff et al., 2012) is highly immunoreactive to antisera raised against DCO (magenta), a catalytic subunit of protein kinase A (PKA). The immunoreactivity pattern of a novel neuropil (arrowheads) to the same antibody in two evolutionarily distinct species of stomatopod *S. empusa* (**B**) and *N. oerstedii* (***C***) is comparable. (**D**) Immunolabeling with an antibody raised against a different subunit of PKA, RII (amber), also revealed the same neuropil immediately posterior to the CB. (**E-G**) While all neuropils, including those of the CX, are immunoreactive to anti synapsin (blue), only the CBii and the newly discovered neuropil react to antisera raised against glutamate decarboxylase (yellow), the enzyme that catalyzes the creation of the inhibitory neurotransmitter GABA. (**H**) The CBii and the undescribed neuropil reacted very weakly to anti-GABA. This anti-GAD and anti-GABA reactivity pattern is reminiscent of that seen in many insects. In the dragonfly *Aeschna cyanea* (**I**; adapted from Homberg et al., 2018), the superior layer (GU1) of the central body upper/fan-shaped body (CBU) is not GABA-ergic. However, both the inferior layer (GU2) of the central body upper and the central body lower/ellipsoid body (CBL) react strongly to anti-GABA. Scale bars = 50 um in **A**-**H**; 100 um in **I**.

Western Blot data comparing the antigenicity of DCO in *N. oerstedii*, the crayfish *Procambarus clarkii,* the fruit fly *Drosophila melanogaster*, and the tropical house cricket *Gryllodes sigillatus* heads reveal a single band at ∼40 kDa (the expected molecular weight of DCO) for all species, indicating high antibody specificity (Supp. Fig. 2A; see also Wolff and Strausfeld, 2015). In both *N. oerstedii* and *Haptosquilla trispinosa*, the region in which this structure would be found was α-tubulin negative (Supp. Fig. 2B-D). Additionally, in those two species, the neuropil is caudoventral to the CB, whereas in *S. empusa* it is far more caudal– so much so that it is offset from the CB (Fig. 4B).

Because the EB in many insects is also reactive to antibodies raised against the inhibitory neurotransmitter gamma-aminobutyric acid (GABA), immunohistochemistry was performed with antisera raised against glutamate decarboxylase (GAD), the enzyme responsible for catalyzing glutamate to GABA, as well as antisera raised against GABA. Both the lower division of the CB as well as the DCO-labeled neuropil were GAD- and GABA-positive, whereas the upper division of the CB was GAD- and GABA-negative (Fig. 4E-H).

### Organization of the stomatopod lateral complex

Stomatopods possess insect-like LXs, with prominent lateral accessory lobes (LALs) as well as two sets of paired bulbs (BUs) that extend laterally from the CB (Fig. 5; Thoen et al., 2017b). In *N. oerstedii*, the LALs are paired oblong neuropils that reside ventrolateral to the CB (Fig. 5A, B). The two pairs of BUs, which in earlier decapod studies were termed lateral lobes, extend out laterally from the lower division of the CB on both sides (Utting et al., 2000). These structures correspond to the stomatopod LALs and BUs described in Thoen et al. (2017) and are structurally and positionally similar to the LX in insects. Some insect species similarly possess multiple pairs of BUs (Held et al., 2016; Hulse et al., 2021). Because the anatomical arrangement of these structures in stomatopods differs from that in those insects that also possess multiple pairs of bulbs (such as honeybees), and because it is highly difficult to determine which pairs of BU may correspond to each other, here we refer to them as the anterior bulbs (aBU) and the posterior bulbs (pBU) in accordance with their anatomical positions (Fig. 5 F, G; Hensgen et al., 2021; Held et al., 2016; Hulse et al., 2021).

**Figure 5.**
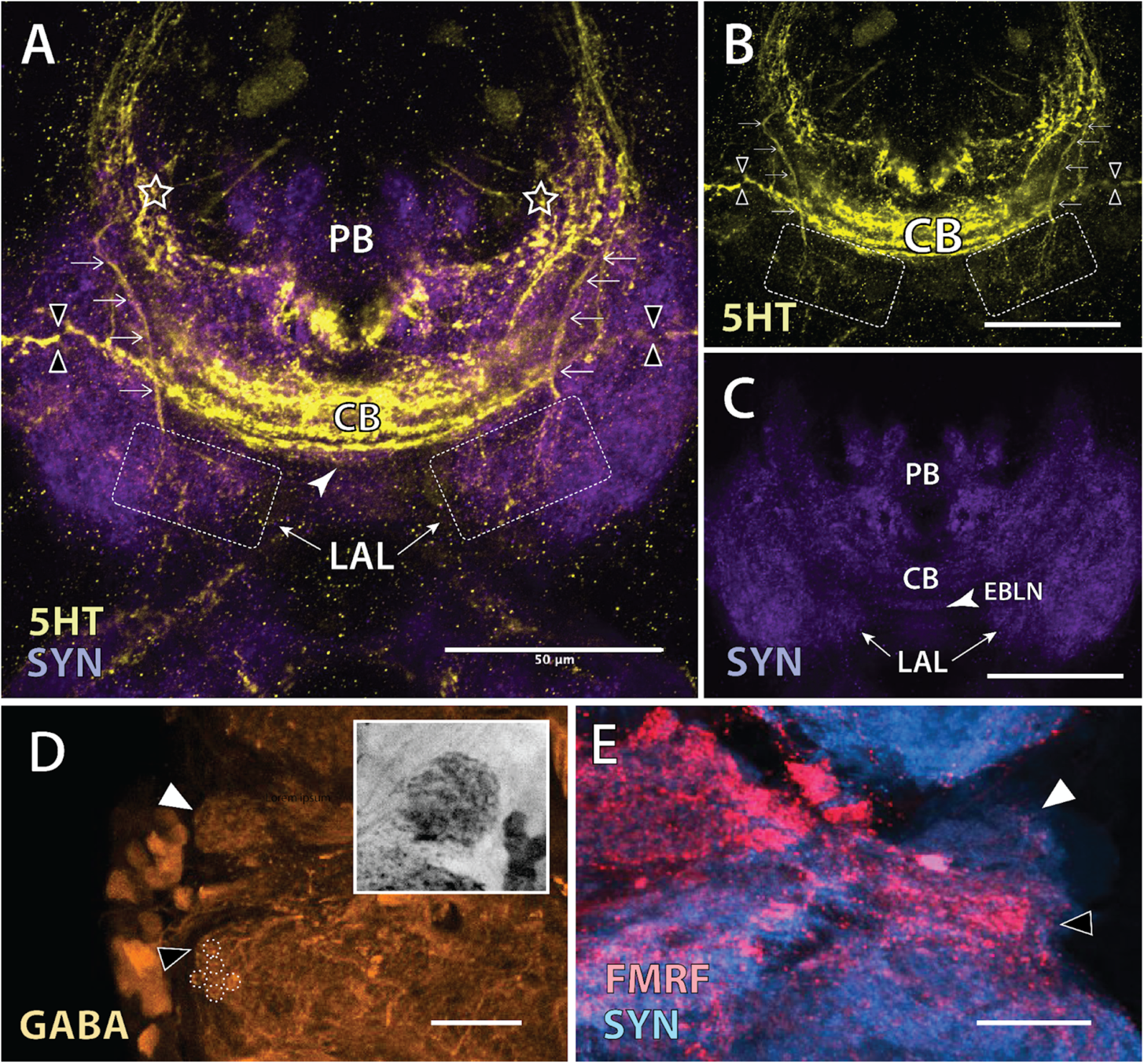
Two sets of neuropils comprise the lateral complex in stomatopods: the lateral accessory lobes (LAL) and the anterior and posterior bulbs (aBU and pBU). (**A**-**C**) The LALs (dotted boxes) are a pair of neuropils lateral to the EBLN. Immunostaining of *N. oerstedii* larval brains with antisera raised against serotonin (5HT; yellow) and synapsin (SYN; purple) reveals connectives (double triangles) between the central body (CB), the EBLN (arrowhead), and the bulbs. It also resolves a pair of bilateral neurons with broad ramifications in the LALs. The axons (arrows) seem to originate from the anterior-most portion of the lateral protocerebrum (stars). The bulbs extend out distally relative to the CB. (**D, E**) The aBU (closed triangles) and pBU (open triangles) are defined by the presence of microglomerular synaptic complexes (examples circled in the pBU) that are strongly immunoreactive to anti-GABA (**D**; orange; grayscale in the inset). (**E**) The two BU can also be differentiated when labeling with antisera raised against FMRFamide (red). The pBU is FMRFamide positive (black triangle), whereas the aBU is not (white triangle). Scale bars = 50 um.

Characteristic features of insect medial and lateral bulbs include large microglomerular structures, which originate from large pre-synaptic sites between neurons that project from the anterior optic tubercle (AOTu; a region thus far undescribed in stomatopods) to GABAergic tangential “ring” neurons with output arborizations in the EB (Mota et al., 2016; Held et al., 2016; von Hadeln et al., 2021). In flies, these microglomeruli are arranged retinotopically (Seelig and Jayaraman, 2013; Hulse et al., 2021). In stomatopods, conspicuous microglomeruli can be resolved with anti-GABA immunolabeling in both the aBU and the pBU (Fig. 5F).

Immunostaining with antisera raised against 5HT reveals several intriguing neuronal axons (Fig. 5C-E). First, a bilateral pair of neurons extends from the anterior lateral protocerebrum and ramifies in the lateral accessory lobes. While neither an anterior nor a posterior optic tubercle has been described thus far in stomatopod crustaceans, the axons seem to originate in an area between the eyestalk, at the base of the protocerebral tract, and the rest of the midbrain. The region is located on the ventral side of the stomatopod brain, which in insects would be the anterior side given the brain rotation. There are also discernible neuronal axons visible with anti-5HT labeling that extend out laterally from the EBLN that terminate in the aBUs. In contrast, immunolabeling with anti-FMRFamide resolves neurons from CBii that ramify in the pBU (Fig. 5F).

### Central complex organization in developing stomatopod crustaceans

Major developmental transitions, such as arthropod metamorphosis, have deep implications for post-embryonic neurogenesis. As they pass through their often dramatically different life stages, animals possess distinct behavioral and morphological adaptations best suited to the ecological niches which they currently occupy. Consequently, nervous systems must also reorganize to effectively meet these demands.

As in many other marine crustaceans, the mantis shrimp development involves a complex ecological shift from free-swimming larval stages among the plankton to benthic juvenile-adult forms. After hatching, the mantis shrimp *N. oerstedii* proceeds through pro-pelagic stages (Stages I-III) in which they are thigmokinetic and negatively phototactic, then molt into stages that are planktonic and free-swimming (Stages IV-VII) (Manning and Provenzano, 1978; Provenzano and Manning, 1978). Larval development ends upon molting into the penultimate postlarval stage. Immunostaining of larval *N. oerstedii* midbrains (Stages III, IV, and VI) revealed a fairly adult-like CX. The PB in Stage II larval *N. oerstedii* appears to possess the same gross morphology as the PB in adults, in which the body of the PB crosses the midline and has two arm-like extensions that curve antero-ventrally (Supp Fig. 1C). Immunostaining with antisera raised against 5HT highlights a CB that spans nearly the entire width of the midbrain. The CB curves anteriorly and extends towards the lateral protocerebrum. As in adults, this structure is divided into an upper (CBi) and lower layer (CBii).

Like adults, *N. oerstedii* larvae possess an EBLN. EBLNs in larvae are strongly immunoreactive to antisera raised against DCO and RII (Supp. Fig. 2D). In fact, all CX neuropils we describe in adult stomatopods are present in *N. oerstedii* larvae, suggesting that these major neuropils form during embryonic development and are important for all developmental stages. This is in contrast to the heterochronic development of some of their sensory neuropils (Supp. Fig. 3). The gross structure of the CB is thin and curved anterolaterally, and spans nearly the entire width of the brain (Supp. Fig. 4A-C).

Larval stomatopods possess a lateral protocerebrum with multiple lobed structures which are not seen in adults (Supp. Fig. 4). In *N. oerstedii* Stage III and Stage IV larvae, these lateral protocerebral lobes (LPL) are strongly reactive to antisera raised against RII. Subsequently, in stage VI, anti-RII immunoreactivity has declined significantly. In contrast, this region in adult stomatopods is diffuse and relatively undefined.By Stage VI, anti-RII immunoreactivity has declined substantially in the LPL. The subunits of this structure can be further differentiated by both anti-5HT and anti-FMRF immunolabeling. When immunolabeled with the former, the posterior-most lobe is labeled very prominently. An axon extends from this lobe down to the ventral nerve cord. The same posterior lobe, as well as the two anterior-most lobes, are additionally immunoreactive to anti-GAD and anti-RII. Equivalent structures are also immunoreactive to anti-RII.

## Discussion

### Considering malacostracan crustacean CXs

Across hexapods and malacostraca, modular midline neuropils can be found with a number of shared characteristics. In all insects analyzed to date, the central complex (CX) comprises an elongated protocerebral bridge and a bipartite central body, consisting of an upper division (or the fan-shaped body) and a lower division (the ellipsoid body). Pterygote (winged) insects have the addition of the paired noduli. Most current understanding of the CX originates from such insect species as locusts, crickets, flies, cockroaches, and butterflies. Recent studies of these insects have implicated the CX in sensory integration, motor coordination, spatial orientation. However, much less is known about CXs in other arthropods.

Within malacostracan crustacean CXs, shared features with insects can be found in the 8/16-fold columnar neuroarchitecture, the presence of an elongated midline neuropil (the PB), a dense and prominent mindline neuropil (the CB), the four pairs of fiber bundles that connect the PB and CB, and connections with the lateral complex (Utting et al., 2000; Homberg et al., 2008). However, neuroanatomical studies of the cerebral ganglia have primarily been documented in the orders Decapoda and Isopoda (Homberg, 2008; Sandeman et al., 1992; Holmgren, 1916; Thompson et al., 1994; Loesel et al., 2002; Utting et al., 2000). In limited previous studies, the upper and lower divisions of the large, midline-spanning neuropil found across Crustacea have been considered to correspond to the insect FB and EB, respectively (Homberg, 2008; Utting et al., 2000; Theon et al., 2017b).

Our data suggest that stomatopod crustaceans possess an additional central complex neuropil exhibiting phenotypic similarity to insect EB in that it is immunoreactive to antisera raised against DCO and GAD. The lower division of the CB is also weakly GAD-positive. There are two possible conclusions that can be inferred from these findings. One is that the two GAD-positive regions are a single compound structure, where the previously undescribed bulbous neuropil of stomatopods is an elaboration of the lower division of the CB. However, studies have shown that some tangential neurons in the FB are GABA-ergic, and multiple dicondylic insects, including visual predators such as praying mantis and dragonflies, exhibit positive GABA immunostaining in the lower division of their FBs (Fig. 4I; Homberg et al., 2018). Thus, the second possibility is that the phenotypic similarity of this previously undescribed neuropil to the insect EB implies that what was once considered to be a bi-layered CB is a bi-layered neuropil analogous to the insect FB. This would suggest that the prominent and bi-layered midline neuropil in stomatopods may be analogous to the insect FB, especially when considered in the context of the discovery of the EBLN (Fig. 6). Furthermore, it implies that the prominent midline neuropils in other malacostracan crustaceans, such as decapods and isopods, are analogous to FBs, and not CBs. The terms homologous and analogous respectively imply similar evolutionary origins, or the lack thereof. Here, we refer to structures as analogous not to rule out potential homology, but because it is the more conservative term.

**Figure 6.**
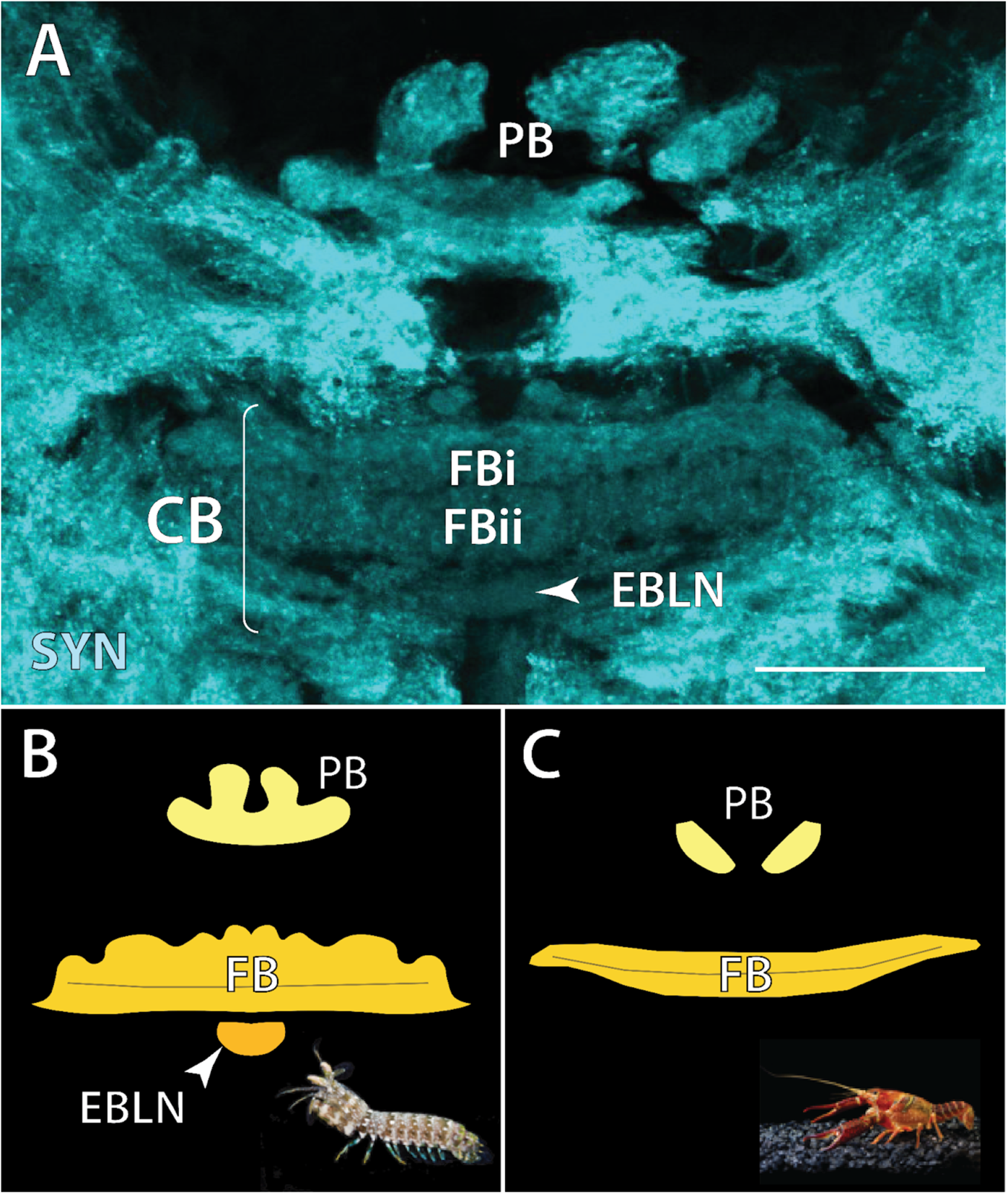
Proposed identification of malacostracan crustacean central complexes. (**A, B**) In stomatopods, if the newly described neuropil (EBLN; arrowhead) corresponds to the insect ellipsoid body, the prominent, bi-layered midline neuropil (FBi and FBii) is likely analogous to the insect fan-shaped body. It follows that the equivalent midline neuropil in other crustaceans, such as that seen in this diagram the crayfish *Procambarus clarkii* CX (**C**) is also a FB. Thus, the central complex in stomatopods consists of an elaborate PB, a bi-layered FB, and an EBLN. The latter two would comprise the central body (CB). In other malacostracan crustaceans, only a PB and FB have been described.

As noted in an earlier publication, stomatopods seem to have CX structures that are unique among malacostracan crustaceans (Thoen et al., 2017b). An example is the presence of the paired NO, two sphere-shaped units that in insects appear to be involved in relaying self-motion information to other CX neuropils (Thoen et al., 2017b; Lu et al., 2021; Lyu et al., 2021; Stone et al., 2017). However, we were unable to identify any clear NO structures in our present study. Thus, we conclude that stomatopod CXs consist of an elaborate PB, a FB, and an EBLN, whereas other malacostracan crustaceans may only possess a simple, bilateral PB and a FB. The exact relationship between insects and crustaceans has been a matter of debate for decades– as has been CX homology, or the lack thereof, among arthropods in general (Strausfeld et al., 2016; Oakley et al., 2013; Osorio et al., 1995; Thoen et al., 2017b; Strausfeld, 1998; Schwentner et al., 2017; Richter, 2002; Loesel, 2004). Due to the ancient evolutionary radiation of Stomatopoda, the presence of the EBLN in stomatopods contributes to the continued studies of neurophylogeny, and generates numerous questions about its potential function. Some of these questions will be discussed in a later section.

Furthermore, assuming crustacean EBs (or EBLNs) play similar functional roles as EBs in insects, it is likely that head direction circuits are present in some malacostracan crustaceans besides stomatopods. The head direction circuit in insect EBs seems to be extremely conserved. Connectomic studies in flies and bees indicate that these animals share similarities in the numbers, types, and projection patterns of CX columnar cells projecting to the EB, whereas these anatomical features appear to be more variable in the FB and NO (Hulse et al., 2021; Sayre et al., 2021). Thus, comparative connectomic and functional studies are necessary to gain a comprehensive understanding of the unusual stomatopod CX.

### An adult-like central complex neuropil in pro-pelagic larvae

CX development has been described in both hemimetabolous and holometabolous insects (Boyan and Liu, 2016; Koniszewski et al., 2016; Young and Armstrong, 2010). Broadly, the development of the final form of the CX coincides with the manifestation of compound eyes and walking legs, although each of the component neuropils is added at distinctive time points (Pfeiffer and Homberg, 2014). In insects like grasshoppers and cockroaches that undergo incomplete metamorphosis, also known as hemimetaboly, the CX is present in an adult-like form in the earliest stages (Koniszewski et al., 2016). In contrast, insects that go through complete metamorphosis may completely lack some neuropils in the larval stages. For example, in the fruit fly *D. melanogaster*, the PB is present in the first instar but the FB does not appear until the third instar (Young and Armstrong, 2010). The EB and NO appear later still, during pupation.

The developmental program of crustaceans is suspected to be more similar to that of hemimetabolous insects than of holometabolous insects, based on the development of compound eyes in a few decapod species (Harzsch et al., 1999). As in hemimetabolous insects, the basic body plan is produced embryonically. Thus, one might expect their CX development to be similar. However, change in malacostracan crustacean central nervous systems past hatching is far more prevalent (Harzsch and Dawirs, 1996; Harzsch et al., 1998; Schmidt and Harzsch, 1999; Sandeman et al., 2011; Cayre et al., 2007; Schmidt, 2014). Futhermore, post-embryonic life for most marine crustaceans involves an often complex ecological shift from free-swimming larval stages among the plankton to benthic juvenile-adults. During this time, there can be multiple molts, where the animal increases in size and sometimes adds more segments and appendages (Haug, 2020). In most stomatopod species, mothers carry and brood gelatinous, yolky egg masses until they hatch. Once the young emerge, they remain in the burrow for one to three pro-pelagic stage(s) and continue to subsist on their yolk (Dingle, 1969; Schram et al., 2013). The pelagic stages begin when the larvae leave the burrow, enter the water column, and become positively phototactic. At this point, the yolk has been consumed and the larvae become predatory much like adults. Although larval raptorial appendages lack the morphological specializations found in adults, they utilize the same latch-mediated spring actuation mechanism to generate high-acceleration strikes (Harrison et al., 2021). After several more molts, the pelagic larvae metamorphose into the post-larval stage, which closely resembles adults and possesses the full complement of stomatopod appendages. Are stomatopod CXs remodeled post-embryonically to reflect these dynamic changes in ecology, behavior, and anatomy?

Critically, newly hatched stomatopod larvae possess the full complement of CX neuropils (Fig. 7). They have a fully formed PB, a bi-layered FB, and an EBLN. The adult-like structure of their CXs suggests that these major neuropils were formed during embryogenesis. However, the FB does not achieve its full adult neuroarchitecture in larvae. The larval FB is much wider relative to the rest of the brain than it is in adults, and its gross structure is thin and curved, as opposed to the fan shape for which adult FB are named. Larval FBs are also highly laminated, though less so than in adults. The relatively complex organization of the larval CX compared to the rest of the brain suggests that the fundamental functions of the CX are critical throughout all life stages.

**Figure 7.**
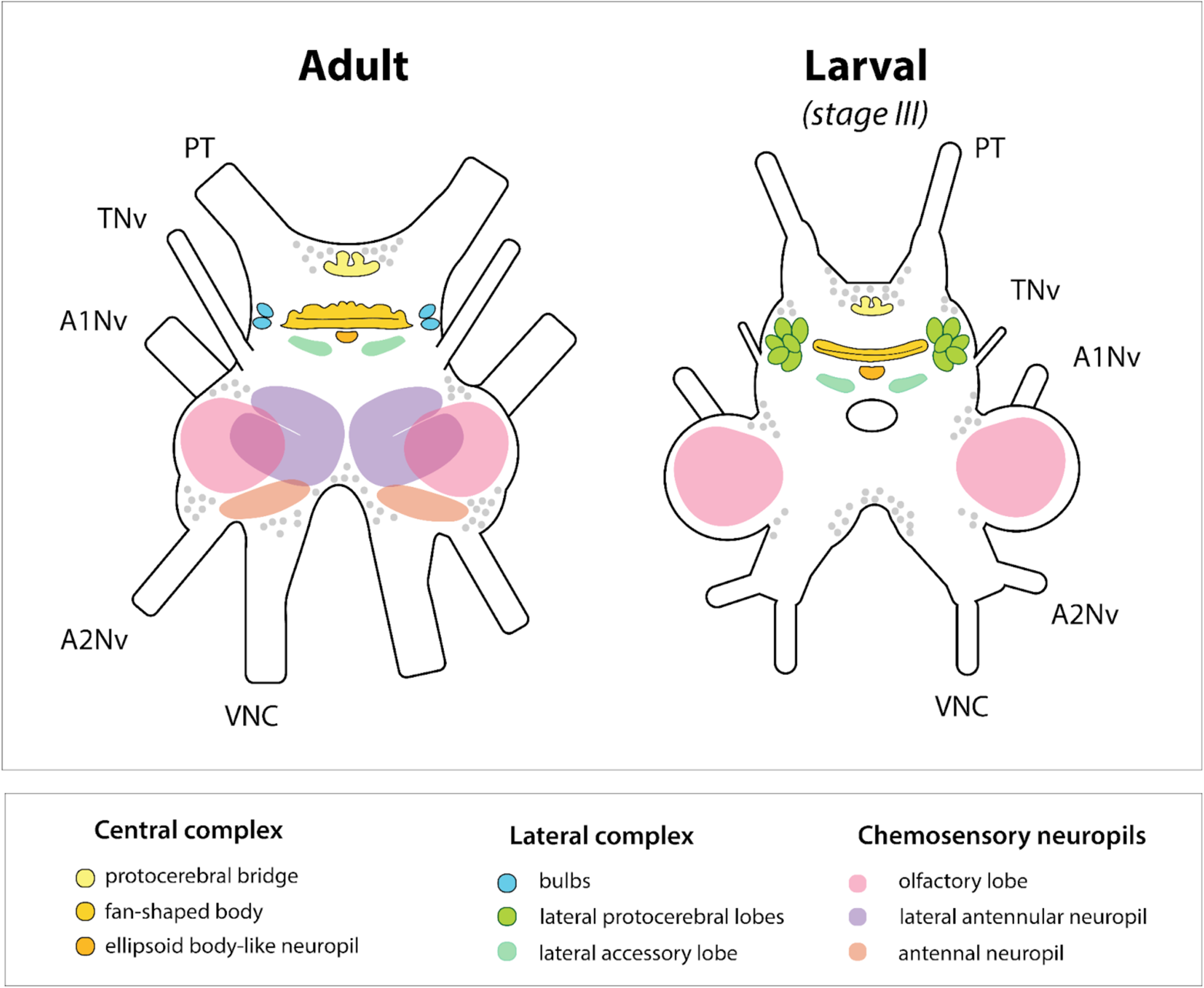
Schematic diagram to compare organization of select neuropils in adult and Stage III larval stomatopod central brains. Larval stomatopods possess all central complex neuropils found in adults (the protocerebral bridge, fan shaped body, and ellipsoid-body like neuropil). However, they have lateral protocerebral lobes where the bulbs would be found in adults. Furthermore, early stage larvae lack lateral antennular neuropils and antennal neuropils. Protocerebral tract (PT); tegumentary nerves (TN); antennular (antenna I) nerve (A1Nv); antennal (antenna II) nerve (A2Nv); and the ventral nerve cord (VNC). Not to scale.

The timing of stomatopod post-embryonic nervous system development is overall poorly studied, with the exception of their unusual eye development. Stomatopod larvae possess apposition compound eyes typical of many crustacean larvae, without the midband specializations that characterize adult eyes. Instead of modifying the existing larval visual system, an adult retina and corresponding optic lobes emerge adjacent to the larval retina and visual processing centers just prior to metamorphosis (Feller et al., 2015; Lin and Cronin, 2018). In other words, late-stage stomatopod larvae and post-larvae possess two sets of visual systems, giving rise to questions about downstream sensory processing. Based on our findings, early stage larval stomatopods also possess unusual lobular structures, which here we call lateral protocerebral lobes (LPL), in a region that is more diffuse and unorganized in adults. Their similarity to the paired bulbs in both shape and location (although not number) in adult stomatopods suggests their potential to be modular visual relay centers whose organization (or lack thereof) corresponds with the changes in visual systems during development.

### A putative neural substrate for navigation and orientation

Across numerous insect clades, including Hymenoptera, Diptera, and Orthoptera, the central complex, in conjunction with the lateral complex, has been shown to be a major component of the neural circuitry underlying navigation and orientation behavior, acting as both a sensory integrator and premotor center. CX neuroarchitecture is characteristically regular, with 16-18 vertical columns that are intersected by horizontal layers (Williams, 1975; Wolff et al., 2015). Columnar cells connect corresponding vertical modules of the PB to the CB; in several insects the modules have been shown to be neurotopic maps of the sensory surround. Each of the horizontal layers is innervated by tangential cells that provide information from other parts of the brain (Pfeiffer and Homberg, 2014). Lesion studies in insects suggest that these layers, especially within the FB, are important for coding visual cue information (Wang et al., 2008, Pan et al., 2009). Sensory inputs are indirect and predominantly visual, although CX neurons have also been implicated in the processing of olfactory, gustatory, and mechanosensory information (Bouhouche et al., 1993; Dus et al., 2011; Okubo et al., 2020).

One of the best-studied functions of the CX is the coding of celestial compass cues for spatial orientation and long-range navigation. In insects such as dung beetles, moths, locusts, and flies the CX uses the sky polarization pattern and positions of the sun or moon to inform orientation and navigation behavior (Beetz and Jundi, 2018; Dacke et al., 2004; Heinze and Homberg 2007; Homberg et al., 2011; Sauman et al., 2005; Honkanen et al., 2019; Giraldo et al., 2018). In insects such as the locust *S. gregaria* and the fruit fly *D. melanogaster,* polarization information travels along the anterior visual pathway via the anterior optic tubercle, through the bulbs, ultimately reaching the EB (Hardcastle et al., 2021; Homberg et al., 2011; Pfeiffer et al., 2005). Furthermore, a ring attractor network has been identified in the PB of flies that encodes a given fly’s heading (Kim et al., 2017; Seelig and Jayaraman, 2013; Turner-Evans et al., 2020).

The orientation and navigational strategies utilized by stomatopod crustaceans are highly comparable to those used by model insect systems. The stomatopod *N. oerstedii* is the only fully aquatic animal known to path integrate (Patel and Cronin, 2020a). This behavior is informed by the sun’s position, overhead polarization patterns, and internal cues. The use of celestial compass cues in particular is strikingly similar to how many terrestrial arthropods navigate. Results from this work and previous studies provide a preliminary framework for understanding stomatopod neural circuitry, particularly in those regions underlying polarization processing and navigation. For example, the presence of an EBLN poses some fascinating questions. Does it serve a similar role to the ellipsoid body in other arthropods? Specifically, does it encode and maintain an internal representation of body orientation or heading? The stomatopod FB is also extensively laminated. Could this elaborate CB neuroarchitecture contribute to evaluating self-motion cues when navigating the benthos, or be used to note environmental visual features salient to their behaviors? We also describe anterior and posterior pairs of bulbs with distinct microglomerular structures. Similar microglomerular synaptic clusters in the BUs have been described in insects such as *D. melanogaster*, the moth *Manduca sexta*, the locust *Schistocerca gregaria,* and the honeybee *Apis mellifera*– all insects that use the sky polarization pattern to navigate (Hanesch et al., 1989; Seelig and Jayaram, 2013; Homberg et al., 1990; Träger et al., 2008; Held et al., 2016; Mota et al., 2016). Histological studies in honeybees indicate that GABAergic tangential neurons connect the BU to the EB (Held et al., 2016; Mota et al., 2016). These connections are retinotopic, and part of the parallel pathways that relay sky-compass information, specifically polarization information, from the dorsal rim area (DRA) of the compound eye to the central complex. Although an analogous dorsal rim area has not been definitively identified in stomatopod crustaceans, molecular studies suggest that the ommatidia in the dorsal margin of *N. oerstedii* eyes, which have visual axes that generally face upwards, have a unique opsin expression pattern (Porter et al., 2020).

## Conclusion

In this study, we detail key characteristics of the stomatopod central complex, including an ellipsoid body-like neuropil, lamination of a putative fan-shaped body (or what was previously considered the stomatopod central body), and microglomerular complexes within the bulbs. The neural basis of insect navigation features the central complex as its control center. Neurobiological correlates of current and desired headings have both been described in insect CXs, which have also long been established as a premotor center. While many regions of stomatopod brains are still undefined, this work lends support to the possibility of analogous navigationally relevant neural circuitry in stomatopods. Whether or not these similarities are products of convergent evolution or are evidence of homology is still up for discussion. Regardless, that stomatopod neurobiology demands further investigation is unquestionable. Their intricate sensory abilities, predatory prowess, and navigational abilities make them an intriguing neuroethological subject– one with strong potential to help understand the evolutionary origins of arthropod navigation and orientation.

## Acknowledgements

This research was supported by grants from the Air Force Office of Scientific Research under grant no. FA9550-18-1-0278 and the University of Maryland Baltimore County (SR18CRON). The authors would like to thank Judy Ching-Wen Wang and Amy Streets for their generous assistance in collecting larval stomatopods. The authors would also like to extend their gratitude to Tagide de Carvalho and the UMBC Keith R. Porter Imaging facility for assistance in confocal imaging.

## References

Beetz MJ, El Jundi B. 2018. Insect Orientation: Stay on Course with the Sun. Curr Biol. doi:10.1016/j.cub.2018.07.032

Bouhouche A, Vaysse G, Corbiegre M. 1993. Immunocytochemical and Learning Studies of a Drosophila Melanogaster Neurological Mutant, No-BridgeKS49 as an Approach to the Possible Role of the Central Complex. J Neurogenet 9:105–121. doi:10.3109/01677069309083453

Boyan GS, Liu Y. 2016. Development of the Neurochemical Architecture of the Central Complex. Front Behav Neurosci 10:167. doi:10.3389/fnbeh.2016.00167

Byrne M, Dacke M, Nordström P, Scholtz C, Warrant E. 2003. Visual cues used by ball-rolling dung beetles for orientation. J Comp Physiol A Neuroethol Sens Neural Behav Physiol 189:411–418. doi:10.1007/s00359-003-0415-1

Cayre M, Scotto-Lomassese S, Malaterre J, Strambi C, Strambi A. 2007. Understanding the regulation and function of adult neurogenesis: contribution from an insect model, the house cricket. Chem Senses 32:385–395. doi:10.1093/chemse/bjm010

Chiou T-H, Kleinlogel S, Cronin T, Caldwell R, Loeffler B, Siddiqi A, Goldizen A, Marshall J. 2008. Circular polarization vision in a stomatopod crustacean. Curr Biol 18:429–434. doi:10.1016/j.cub.2008.02.066

Crittenden JR, Skoulakis EM, Han KA, Kalderon D, Davis RL. 1998. Tripartite mushroom body architecture revealed by antigenic markers. Learn Mem 5:38–51.

Cronin TW, Johnsen S, Justin Marshall N, Warrant EJ. 2014. Visual Ecology. Princeton University Press.

Cronin TW, Marshall J. 2001. Parallel processing and image analysis in the eyes of mantis shrimps. Biol Bull 200:177–183. doi:10.2307/1543312

Cronin TW, Marshall NJ. 1989a. Multiple spectral classes of photoreceptors in the retinas of gonodactyloid stomatopod crustaceans. Journal of Comparative Physiology A 166:261–275. doi:10.1007/BF00193471

Cronin TW, Marshall NJ. 1989b. A retina with at least ten spectral types of photoreceptors in a mantis shrimp. Nature 339:137–140. doi:10.1038/339137a0

Cronin TW, Shashar N, Caldwell RL, Marshall J, Cheroske AG, Chiou T-H. 2003. Polarization vision and its role in biological signaling. Integr Comp Biol 43:549–558. doi:10.1093/icb/43.4.549

Dacke M, Byrne MJ, Baird E, Scholtz CH, Warrant EJ. 2011. How dim is dim? Precision of the celestial compass in moonlight and sunlight. Philos Trans R Soc Lond B Biol Sci 366:697–702. doi:10.1098/rstb.2010.0191

Dacke M, Byrne MJ, Scholtz CH, Warrant EJ. 2004. Lunar orientation in a beetle. Proc Biol Sci 271:361–365. doi:10.1098/rspb.2003.2594

Dacke M, El Jundi B. 2018. The Dung Beetle Compass. Curr Biol 28:R993–R997. doi:10.1016/j.cub.2018.04.052

Dacke M, Nordström P, Scholtz CH. 2003. Twilight orientation to polarised light in the crepuscular dung beetle Scarabaeus zambesianus. J Exp Biol 206:1535–1543. doi:10.1242/jeb.00289

Donlea JM, Pimentel D, Miesenböck G. 2014. Neuronal machinery of sleep homeostasis in Drosophila. Neuron 81:860–872. doi:10.1016/j.neuron.2013.12.013

Dus M, Min S, Keene AC, Lee GY, Suh GSB. 2011. Taste-independent detection of the caloric content of sugar in Drosophila. Proc Natl Acad Sci U S A 108:11644–11649. doi:10.1073/pnas.1017096108

El Jundi B, Baird E, Byrne MJ, Dacke M. 2019. The brain behind straight-line orientation in dung beetles. J Exp Biol 222:jeb192450–7. doi:10.1242/jeb.192450

El Jundi B, Warrant EJ, Byrne MJ, Khaldy L, Baird E, Smolka J, Dacke M. 2015. Neural coding underlying the cue preference for celestial orientation. Proc Natl Acad Sci U S A 112:11395– 1400. doi:10.1073/pnas.1501272112

Furshpan EJ, Potter DD. 1959. Transmission at the giant motor synapses of the crayfish. J Physiol 145:289–325. doi:10.1113/jphysiol.1959.sp006143

Giraldo YM, Leitch KJ, Ros IG, Warren TL, Weir PT, Dickinson MH. 2018. Sun Navigation Requires Compass Neurons in Drosophila. Curr Biol 28:2845–2852.e4. doi:10.1016/j.cub.2018.07.002

Guo P, Ritzmann RE. 2013. Neural activity in the central complex of the cockroach brain is linked to turning behaviors. J Exp Biol 216:992–1002. doi:10.1242/jeb.080473

Grob R, Fleischmann PN, Grübel K, Wehner R, Rössler W. 2017. The Role of Celestial Compass Information in Cataglyphis Ants during Learning Walks and for Neuroplasticity in the Central Complex and Mushroom Bodies. Front Behav Neurosci 11:226. doi:10.3389/fnbeh.2017.00226

von Hadeln J, Hensgen R, Bockhorst T, Rosner R, Heidasch R, Pegel U, Quintero Pérez M, Homberg U. 2020. Neuroarchitecture of the central complex of the desert locust: Tangential neurons. J Comp Neurol 528:906–934. doi:10.1002/cne.24796

Hanesch U, Fischbach K-F, Heisenberg M. 1989. Neuronal architecture of the central complex in Drosophila melanogaster. Cell Tissue Res 257:343–366. doi:10.1007/BF00261838

Hansen HJ. 1895. Isopoden, Cumaceen und Stomatopoden der Plankton-ExpeditionErgebnisse Plankton-Expedition Der Humboldt-Stiftung. Hardpress Publishing. pp. 1–105.

Hardcastle BJ, Omoto JJ, Kandimalla P, Nguyen B-CM, Keleş MF, Boyd NK, Hartenstein V, Frye MA. 2021. A visual pathway for skylight polarization processing in Drosophila. Elife 10. doi:10.7554/eLife.63225

Harris-Warrick RM, Marder E. 1991. Modulation of neural networks for behavior. Annu Rev Neurosci 14:39–57. doi:10.1146/annurev.ne.14.030191.000351

Harzsch S, Benton J, Dawirs RR, Beltz B. 1999. A new look at embryonic development of the visual system in decapod crustaceans: neuropil formation, neurogenesis, and apoptotic cell death. J Neurobiol 39:294–306.

Harzsch S, Dawirs RR. 1996. Neurogenesis in the developing crab brain: postembryonic generation of neurons persists beyond metamorphosis. J Neurobiol 29:384–398. doi:10.1002/(SICI)1097-4695(199603)29:3<384::AID-NEU9>3.0.CO;2-5

Harzsch S, Miller J, Benton J, Dawirs RR, Beltz B. 1998. Neurogenesis in the thoracic neuromeres of two crustaceans with different types of metamorphic development. J Exp Biol 201 (Pt 17):2465–2479.

Haug JT. 2020. Metamorphosis in crustaceans In: Anger K, Harzsch S, Thiel M, editors. Developmental Biology and Larval Ecology: The Natural History of the Crustacean. Oxford University Press.

Heinze S. 2014. Polarized-Light Processing in Insect Brains: Recent Insights from the Desert Locust, the MonarchPolarized Light and Polarization Vision in Animal Sciences. unknown. pp. 61–111. doi:10.1007/978-3-642-54718-8

Heinze S, Homberg U. 2007. Maplike representation of celestial E-vector orientations in the brain of an insect. Science 315:995–997. doi:10.1126/science.1135531

Heinze S, Narendra A, Cheung A. 2018. Principles of Insect Path Integration. Curr Biol 28:R1043–R1058. doi:10.1016/j.cub.2018.04.058

Heinze S, Reppert SM. 2012. Anatomical basis of sun compass navigation I: the general layout of the monarch butterfly brain. J Comp Neurol 520:1599–1628. doi:10.1002/cne.23054

Held M, Berz A, Hensgen R, Muenz TS, Scholl C, Rössler W, Homberg U, Pfeiffer K. 2016. Microglomerular Synaptic Complexes in the Sky-Compass Network of the Honeybee Connect Parallel Pathways from the Anterior Optic Tubercle to the Central Complex. Front Behav Neurosci 10:186. doi:10.3389/fnbeh.2016.00186

Hensgen R, England L, Homberg U, Pfeiffer K. 2021. Neuroarchitecture of the central complex in the brain of the honeybee: Neuronal cell types. J Comp Neurol 529:159–186. doi:10.1002/cne.24941

Holmgren NF. 1916. Zur vergleichenden anatomie des gehirns: von polychaeten. onychophoren, xiphosuren, arachniden, crustaceen, myriapoden und insekten. Kungliga Svenska Vetenskapsakademiens Handlingar, 56:1–303

Homberg U. 2008. Evolution of the central complex in the arthropod brain with respect to the visual system. Arthropod Struct Dev 37:347–362. doi:10.1016/j.asd.2008.01.008

Homberg U, Heinze S, Pfeiffer K, Kinoshita M, el Jundi B. 2011. Central neural coding of sky polarization in insects. Philos Trans R Soc Lond B Biol Sci 366:680–687. doi:10.1098/rstb.2010.0199

Homberg U, Humberg T-H, Seyfarth J, Bode K, Pérez MQ. 2018. GABA immunostaining in the central complex of dicondylian insects. J Comp Neurol 526:2301–2318. doi:10.1002/cne.24497

Homberg U, Kingan TG, Hildebrand JG. 1990. Distribution of FMRFamide-like immunoreactivity in the brain and suboesophageal ganglion of the sphinx moth Manduca sexta and colocalization with SCPB-, BPP-, and GABA-like immunoreactivity. Cell Tissue Res 259:401–419. doi:10.1007/bf01740767

Honkanen A, Adden A, da Silva Freitas J, Heinze S. 2019. The insect central complex and the neural basis of navigational strategies. J Exp Biol 222:jeb188854–15. doi:10.1242/jeb.188854

Hulse BK, Haberkern H, Franconville R, Turner-Evans DB, Takemura S-Y, Wolff T, Noorman M, Dreher M, Dan C, Parekh R, Hermundstad AM, Rubin GM, Jayaraman V. 2021. A connectome of the Drosophila central complex reveals network motifs suitable for flexible navigation and context-dependent action selection. Elife 10. doi:10.7554/eLife.66039

Ilius M, Wolf R, Heisenberg M. 2007. The central complex of Drosophila melanogaster is involved in flight control: studies on mutants and mosaics of the gene ellipsoid body open. J Neurogenet 21:321–338. doi:10.1080/01677060701693503

Ito K, Shinomiya K, Ito M, Armstrong JD, Boyan G, Hartenstein V, Harzsch S, Heisenberg M, Homberg U, Jenett A, Keshishian H, Restifo LL, Rössler W, Simpson JH, Strausfeld NJ, Strauss R, Vosshall LB, Insect Brain Name Working Group. 2014. A systematic nomenclature for the insect brain. Neuron 81:755–765. doi:10.1016/j.neuron.2013.12.017

Kim SS, Rouault H, Druckmann S, Jayaraman V. 2017. Ring attractor dynamics in the Drosophila central brain. Science 356:849–853. doi:10.1126/science.aal4835

Kleinlogel S, Marshall NJ. 2006. Electrophysiological evidence for linear polarization sensitivity in the compound eyes of the stomatopod crustacean Gonodactylus chiragra. J Exp Biol 209:4262–4272. doi:10.1242/jeb.02499

Kleinlogel S, Marshall NJ. 2005. Photoreceptor projection and termination pattern in the lamina of gonodactyloid stomatopods (mantis shrimp). Cell Tissue Res 321:273–284. doi:10.1007/s00441-005-1118-4

Kleinlogel S, Marshall NJ, Horwood JM, Land MF. 2003. Neuroarchitecture of the color and polarization vision system of the stomatopod Haptosquilla. J Comp Neurol 467:326–342. doi:10.1002/cne.10922

Koniszewski NDB, Kollmann M, Bigham M, Farnworth M, He B, Büscher M, Hütteroth W, Binzer M, Schachtner J, Bucher G. 2016. The insect central complex as model for heterochronic brain development-background, concepts, and tools. Dev Genes Evol 226:209–219. doi:10.1007/s00427-016-0542-7

Labhart T, Meyer EP. 1999. Detectors for polarized skylight in insects: a survey of ommatidial specializations in the dorsal rim area of the compound eye. Microsc Res Tech 47:368–379. doi:10.1002/(SICI)1097-0029(19991215)47:6<368::AID-JEMT2>3.0.CO;2-Q

Lin C, Chou A, Cronin TW. 2020. Optic lobe organization in stomatopod crustacean species possessing different degrees of retinal complexity. J Comp Physiol A Neuroethol Sens Neural Behav Physiol 206:247–258. doi:10.1007/s00359-019-01387-5

Lin C, Cronin TW. 2018. Two visual systems in one eyestalk: The unusual optic lobe metamorphosis in the stomatopod Alima pacifica. Dev Neurobiol 78:3–14. doi:10.1002/dneu.22550

Liu G, Seiler H, Wen A, Zars T, Ito K, Wolf R, Heisenberg M, Liu L. 2006. Distinct memory traces for two visual features in the Drosophila brain. Nature 439:551–556. doi:10.1038/nature04381

Loesel R, Nässel DR, Strausfeld NJ. 2002. Common design in a unique midline neuropil in the brains of arthropods. Arthropod Struct Dev 31:77–91. doi:10.1016/S1467-8039(02)00017-8

Loesel R. 2004. Comparative morphology of central neuropils in the brain of arthropods and its evolutionary and functional implications. Acta Biol Hung 55:39–51. doi:10.1556/ABiol.55.2004.1-4.6

Lu J, Behbahani AH, Hamburg L, Westeinde EA, Dawson PM, Lyu C, Maimon G, Dickinson MH, Druckmann S, Wilson RI. 2022. Transforming representations of movement from body-to world-centric space. Nature 601:98–104. doi:10.1038/s41586-021-04191-x

Lyu C, Abbott LF, Maimon G. 2022. Building an allocentric travelling direction signal via vector computation. Nature 601:92–97. doi:10.1038/s41586-021-04067-0

Manning RB, Provenzano AJ. 1978. Studies on development of stomatopod crustacea I. early larval stages of Gonodactylus oerstedii hansen. Bull Mar Sci 28:467–487.

Marder E. 2012. Neuromodulation of neuronal circuits: back to the future. Neuron 76:1–11. doi:10.1016/j.neuron.2012.09.010

Marshall NJ. 1988. A unique colour and polarization vision system in mantis shrimps. Nature 333:557–560. doi:10.1038/333557a0

Marshall J, Cronin TW, Kleinlogel S. 2007. Stomatopod eye structure and function: a review. Arthropod Struct Dev 36:420–448. doi:10.1016/j.asd.2007.01.006

Marshall NJ, Land MF. 1993. Some optical features of the eyes of stomatopods. Journal of Comparative Physiology A 173:583–594. doi:10.1007/BF00197766

Marshall NJ, Land MF, King CA, Cronin TW. 1991a. The compound eyes of mantis shrimps (Crustacea, Hoplocarida, Stomatopoda). II. Colour pigments in the eyes of stomatopod crustaceans: polychromatic vision by …. Philos Trans R Soc Lond B Biol Sci 334:57–84.

Marshall NJ, Land MF, King CA, Cronin TW, Bone Q. 1991b. The compound eyes of mantis shrimps (Crustacea, Hoplocarida, Stomatopoda). I. Compound eye structure: the detection of polarized light. Philos Trans R Soc Lond B Biol Sci 334:33–56. doi:10.1098/rstb.1991.0096

Martin JP, Guo P, Mu L, Harley CM, Ritzmann RE. 2015. Central-complex control of movement in the freely walking cockroach. Curr Biol 25:2795–2803. doi:10.1016/j.cub.2015.09.044

Mota T, Kreissl S, Carrasco Durán A, Lefer D, Galizia G, Giurfa M. 2016. Synaptic Organization of Microglomerular Clusters in the Lateral and Medial Bulbs of the Honeybee Brain. Front Neuroanat 10:103. doi:10.3389/fnana.2016.00103

Müller M, Wehner R. 1988. Path integration in desert ants, Cataglyphis fortis. Proc Natl Acad Sci U S A 85:5287–5290. doi:10.1073/pnas.85.14.5287

Oakley TH, Wolfe JM, Lindgren AR, Zaharoff AK. 2013. Phylotranscriptomics to bring the understudied into the fold: monophyletic ostracoda, fossil placement, and pancrustacean phylogeny. Mol Biol Evol 30:215–233. doi:10.1093/molbev/mss216

Ofstad TA, Zuker CS, Reiser MB. 2011. Visual place learning in Drosophila melanogaster. Nature 474:204–207. doi:10.1038/nature10131

Okubo TS, Patella P, D’Alessandro I, Wilson RI. 2020. A Neural Network for Wind-Guided Compass Navigation. Neuron 107:924–940.e18. doi:10.1016/j.neuron.2020.06.022

Osorio D, Averof M, Bacon JP. 1995. Arthropod evolution: great brains, beautiful bodies. Trends Ecol Evol 10:449–454. doi:10.1016/s0169-5347(00)89178-8

Pan X, Bourland WA, Song W. 2013. Protargol synthesis: an in-house protocol. J Eukaryot Microbiol 60:609–614. doi:10.1111/jeu.12067

Pan Y, Zhou Y, Guo C, Gong H, Gong Z, Liu L. 2009. Differential roles of the fan-shaped body and the ellipsoid body in Drosophila visual pattern memory. Learn Mem. 16:289–295. doi:10.1101/lm.1331809

Patel RN, Cronin TW. 2020a. Mantis Shrimp Navigate Home Using Celestial and Idiothetic Path Integration. Curr Biol 30:1981–1987.e3. doi:10.1016/j.cub.2020.03.023

Patel RN, Cronin TW. 2020b. Path integration error and adaptable search behaviors in a mantis shrimp. J Exp Biol 223. doi:10.1242/jeb.224618

Patel RN, Cronin TW. 2020c. Landmark navigation in a mantis shrimp. Proc Biol Sci 287:20201898. doi:10.1098/rspb.2020.1898

Patel RN, Khil V, Abdurahmonova L, Driscoll H, Patel S, Pettyjohn-Robin O, Shah A, Goldwasser T, Sparklin B, Cronin TW. 2021. Mantis shrimp identify an object by its shape rather than its color during visual recognition. J Exp Biol 228. doi:10.1242/jeb.242256

Pegel U, Pfeiffer K, Zittrell F, Scholtyssek C, Homberg U. 2019. Two Compasses in the Central Complex of the Locust Brain. J Neurosci 39:3070–3080. doi:10.1523/JNEUROSCI.0940-18.2019

Phillips-Portillo J. 2012. The central complex of the flesh fly, Neobellieria bullata: recordings and morphologies of protocerebral inputs and small-field neurons. J Comp Neurol 520:3088–3104. doi:10.1002/cne.23134

Pfeiffer K, Homberg U. 2014. Organization and functional roles of the central complex in the insect brain. Annu Rev Entomol 59:165–184. doi:10.1146/annurev-ento-011613-162031

Pfeiffer K, Kinoshita M. 2012. Segregation of visual inputs from different regions of the compound eye in two parallel pathways through the anterior optic tubercle of the bumblebee (Bombus ignitus). J Comp Neurol 520:212–229. doi:10.1002/cne.22776

Pfeiffer K, Kinoshita M, Homberg U. 2005. Polarization-sensitive and light-sensitive neurons in two parallel pathways passing through the anterior optic tubercle in the locust brain. J Neurophysiol 94:3903–3915. doi:10.1152/jn.00276.2005

Porter ML, Awata H, Bok MJ, Cronin TW. 2020. Exceptional diversity of opsin expression patterns in Neogonodactylus oerstedii (Stomatopoda) retinas. Proc Natl Acad Sci U S A 117:8948–8957. doi:10.1073/pnas.1917303117

Porter ML, Zhang Y, Desai S, Caldwell RL, Cronin TW. 2010. Evolution of anatomical and physiological specialization in the compound eyes of stomatopod crustaceans. J Exp Biol 213:3473–3486. doi:10.1242/jeb.046508

Popov AV, Peresleni AI, Ozerskii PV, Shchekanov EE, Savvateeva-Popova EV. 2003. On the role of the protocerebral bridge in the central complex of Drosophila melanogaster brain in control of courtship behavior and sound production. J Evol Biochem Physiol 39:655–666. doi:10.1023/b:joey.0000023486.75353.f8

Provenzano AJ Jr, Manning RB. 1978. Studies on Development of Stomatopod Crustacea II. The Later Larval Stages of Gonodactylus Oerstedii Hansen Reared in the Laboratory. Bull Mar Sci 28:297–315.

Richter S. 2002. The Tetraconata concept: hexapod-crustacean relationships and the phylogeny of Crustacea. Org Divers Evol 2:217–237. doi:10.1078/1439-6092-00048

Roberts NW, Porter ML, Cronin TW. 2011. The molecular basis of mechanisms underlying polarization vision. Philos Trans R Soc Lond B Biol Sci 366:627–637. doi:10.1098/rstb.2010.0206

Rossel S, Wehner R. 1984. Celestial orientation in bees: the use of spectral cues. Journal of Comparative Physiology A 155:605–613. doi:10.1007/BF00610846

Sakai T, Kitamoto T. 2006. Differential roles of two major brain structures, mushroom bodies and central complex, for Drosophila male courtship behavior. J Neurobiol 66:821–834. doi:10.1002/neu.20262

Sandeman DC, Bazin F, Beltz BS. 2011. Adult neurogenesis: examples from the decapod crustaceans and comparisons with mammals. Arthropod Struct Dev 40:258–275. doi:10.1016/j.asd.2011.03.001

Sandeman D, Sandeman R, Derby C, Schmidt M. 1992. Morphology of the Brain of Crayfish, Crabs, and Spiny Lobsters: A Common Nomenclature for Homologous Structures. Biol Bull 183:304–326. doi:10.2307/1542217

Sauman I, Briscoe AD, Zhu H, Shi D, Froy O, Stalleicken J, Yuan Q, Casselman A, Reppert SM. 2005. Connecting the navigational clock to sun compass input in monarch butterfly brain. Neuron 46:457–467. doi:10.1016/j.neuron.2005.03.014

Schmidt M. 2014. Adult neurogenesis in crustaceans. The Natural history of crustacea 3:175–206.

Schmidt M, Harzsch S. 1999. Comparative Analysis of Neurogenesis in the Central Olfactory Pathway of Adult Decapod Crustaceans by In Vivo BrdU Labeling. Biol Bull 196:127–136. doi:10.2307/1542558

Seelig JD, Jayaraman V. 2013. Feature detection and orientation tuning in the Drosophila central complex. Nature 503:262–266. doi:10.1038/nature12601

Skoulakis EM, Kalderon D, Davis RL. 1993. Preferential expression in mushroom bodies of the catalytic subunit of protein kinase A and its role in learning and memory. Neuron 11:197–208. doi:10.1016/0896-6273(93)90178-t

Stollewerk A, Simpson P. 2005. Evolution of early development of the nervous system: a comparison between arthropods. Bioessays 27:874–883. doi:10.1002/bies.20276

Stone T, Webb B, Adden A, Weddig NB, Honkanen A, Templin R, Wcislo W, Scimeca L, Warrant E, Heinze S. 2017. An Anatomically Constrained Model for Path Integration in the Bee Brain. Curr Biol 27:3069–3085.e11. doi:10.1016/j.cub.2017.08.052

Schwentner M, Combosch DJ, Pakes Nelson J, Giribet G. 2017. A Phylogenomic Solution to the Origin of Insects by Resolving Crustacean-Hexapod Relationships. Curr Biol 27:1818–1824.e5. doi:10.1016/j.cub.2017.05.040

Strausfeld NJ. 1998. Crustacean – Insect Relationships: The Use of Brain Characters to Derive Phylogeny amongst Segmented Invertebrates. Brain Behav Evol 52:186–206. doi:10.1159/000006563

Strausfeld NJ. 2012. Arthropod brains: Evolution, functional elegance, and historical significance. London, England: Belknap Press. doi:10.2307/j.ctv1dp0v2h

Strausfeld NJ, Ma X, Edgecombe GD. 2016. Fossils and the Evolution of the Arthropod Brain. Curr Biol 26:R989–R1000. doi:10.1016/j.cub.2016.09.012

Strauss R. 2002. The central complex and the genetic dissection of locomotor behaviour. Curr Opin Neurobiol 12:633–638. doi:10.1016/s0959-4388(02)00385-9

Thoen HH, Strausfeld NJ, Marshall J. 2017a. Neural organization of afferent pathways from the stomatopod compound eye. J Comp Neurol 525:3010–3030. doi:10.1002/cne.24256

Thoen HH, Marshall J, Wolff GH, Strausfeld NJ. 2017b. Insect-Like Organization of the Stomatopod Central Complex: Functional and Phylogenetic Implications. Front Behav Neurosci 11:12. doi:10.3389/fnbeh.2017.00012

Thompson KS, Zeidler MP, Bacon JP. 1994. Comparative anatomy of serotonin-like immunoreactive neurons in isopods: putative homologues in several species. J Comp Neurol 347:553–569. doi:10.1002/cne.903470407

Träger U, Wagner R, Bausenwein B, Homberg U. 2008. A novel type of microglomerular synaptic complex in the polarization vision pathway of the locust brain. J Comp Neurol 506:288–300. doi:10.1002/cne.21512

Turner-Evans DB, Jayaraman V. 2016. The insect central complex. Curr Biol 26:R453–7. doi:10.1016/j.cub.2016.04.006

Turner-Evans DB, Jensen KT, Ali S, Paterson T, Sheridan A, Ray RP, Wolff T, Lauritzen JS, Rubin GM, Bock DD, Jayaraman V. 2020. The neuroanatomical ultrastructure and function of a biological ring attractor. Neuron 109:1582. doi:10.1016/j.neuron.2021.04.016

Turner-Evans D, Wegener S, Rouault H, Franconville R, Wolff T, Seelig JD, Druckmann S, Jayaraman V. 2017. Angular velocity integration in a fly heading circuit. Elife 6. doi:10.7554/eLife.23496

Utting M, Agricola H, Sandeman R, Sandeman D. 2000. Central complex in the brain of crayfish and its possible homology with that of insects. J Comp Neurol 416:245–261. doi:10.1002/(sici)1096-9861(20000110)416:2<245::aid-cne9>3.0.co;2-a

Van Der Wal C, Ahyong ST, Ho SYW, Lo N. 2017. The evolutionary history of Stomatopoda (Crustacea: Malacostraca) inferred from molecular data. PeerJ 5:e3844. doi:10.7717/peerj.3844

Wang Z, Pan Y, Li W, Jiang H, Chatzimanoli L, CHang J, Gong Z, Liu L. 2008. Visual pattern memory requires foraging function in the central complex of Drosophila. Learn Mem 15:133–142. doi:10.1101/lm.873008

Waterman TH. 1981. Polarization sensitivity In: Autrum H, editor. Handbook of Sensory Physiology. Springer-Verlag. pp. 281–469.

Wehner R. 1976. Polarized-light navigation by insects. Sci Am 235:106–115. doi:10.1038/scientificamerican0776-106

Weir PT, Schnell B, Dickinson MH. 2014. Central complex neurons exhibit behaviorally gated responses to visual motion in Drosophila. J Neurophysiol 111:62–71. doi:10.1152/jn.00593.2013

Wolff G, Harzsch S, Hansson BS, Brown S, Strausfeld N. 2012. Neuronal organization of the hemiellipsoid body of the land hermit crab, Coenobita clypeatus: correspondence with the mushroom body ground pattern. J Comp Neurol 520:2824–2846. doi:10.1002/cne.23059

Wolff T, Iyer NA, Rubin, GM. 2015. Neuroarchitecture and neuroanatomy of the Drosophila central complex: A GAL4-based dissection of protocerebral bridge neurons and circuits. J Comp Neurol 523:997–1037. doi:10.1002/cne.23705

Young JM, Armstrong JD. 2010. Building the central complex in Drosophila: the generation and development of distinct neural subsets. J Comp Neurol 518:1525–1541. doi:10.1002/cne.22285

