## Supplemental Figures and Legends for "Neuroanatomy of stomatopod central complexes offers putative neural substrate for oriented behaviors in crustaceans"

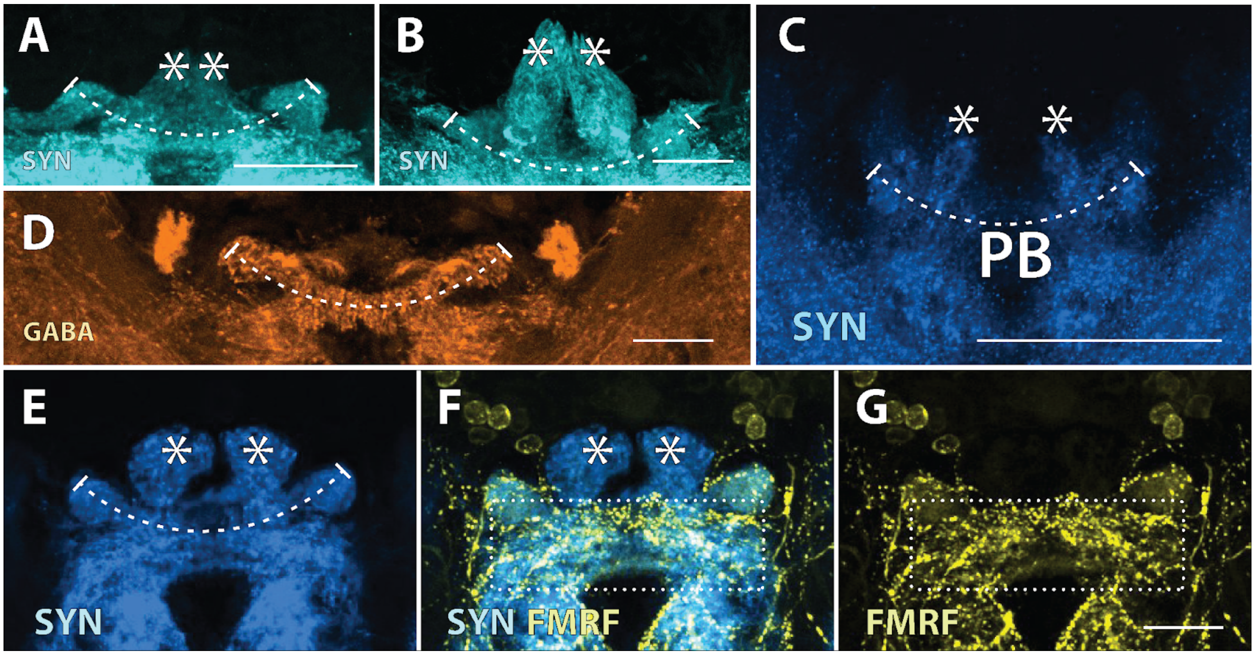


**Supplementary Figure 1**. The stomatopod protocerebral bridge (PB). Its unusual structure consisting of a primary midline portion (dotted section) and two overarching arms (asterisks) is observed in at least three species with anti-synapsin (cyan or blue; SYN) staining, including *Chorisquilla hystrix* (**A**), *Haptosquilla trispinosa* (**B**), and *N. oerstedii* (**C**-**G**). The midline component differentially stains with anti-GABA (orange; **D**) and anti-FMRFamide (yellow; **F, G**). Scale bars = 50 um.

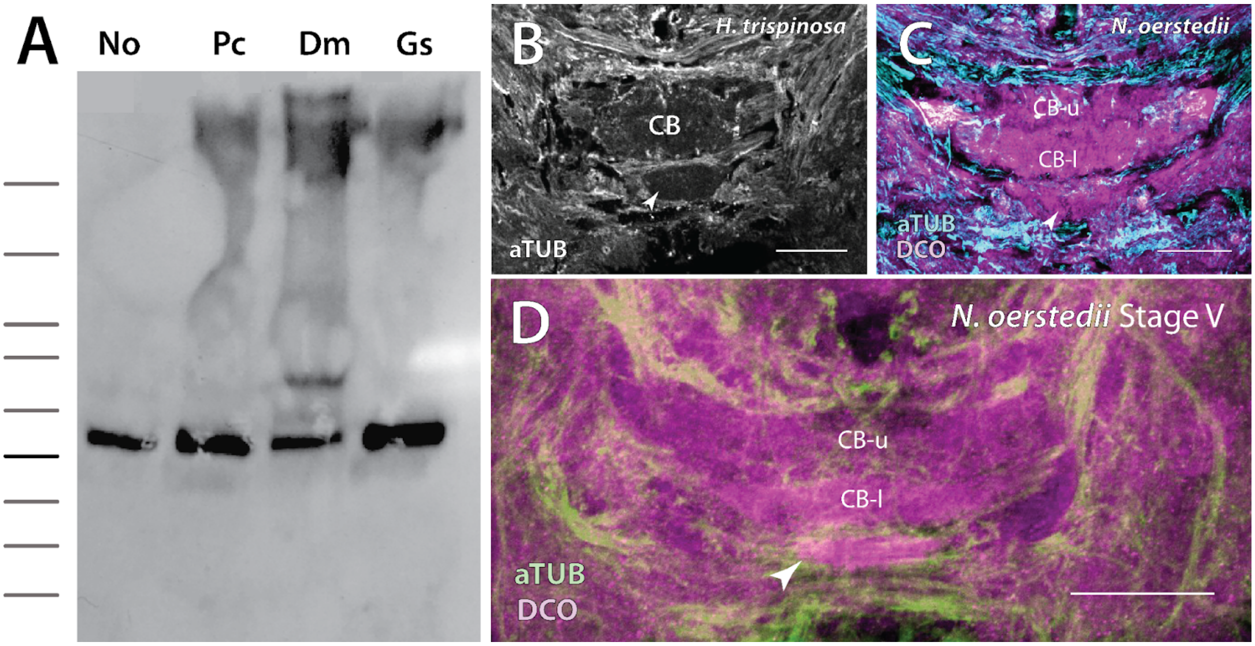


**Supplementary Figure 2**. (**A**) Western Blot of DCO antibody comparing brain tissue of *N. oerstedii* (No), *P. clarkii* (Pc), *D. melanogaster* (Dm), and the cricket *G. sigillatus* (Gs). In each case, a dense band at the expected molecular weight for DCO (~40kDa) is obvious. (**B**) The ellipsoid body-like neuropil is unlikely a cross section of a nerve tract due to its lack of immunoreactivity to anti-alpha tubulin (grayscale; **C**) in *Haptosquilla trispinosa*. Double immunolabeling of anti tubulin (cyan, **C**; green, **D**) with anti-DCO (magenta) in two life stages of *N. oerstedii* show similar results.  Scale bars = 50 um.

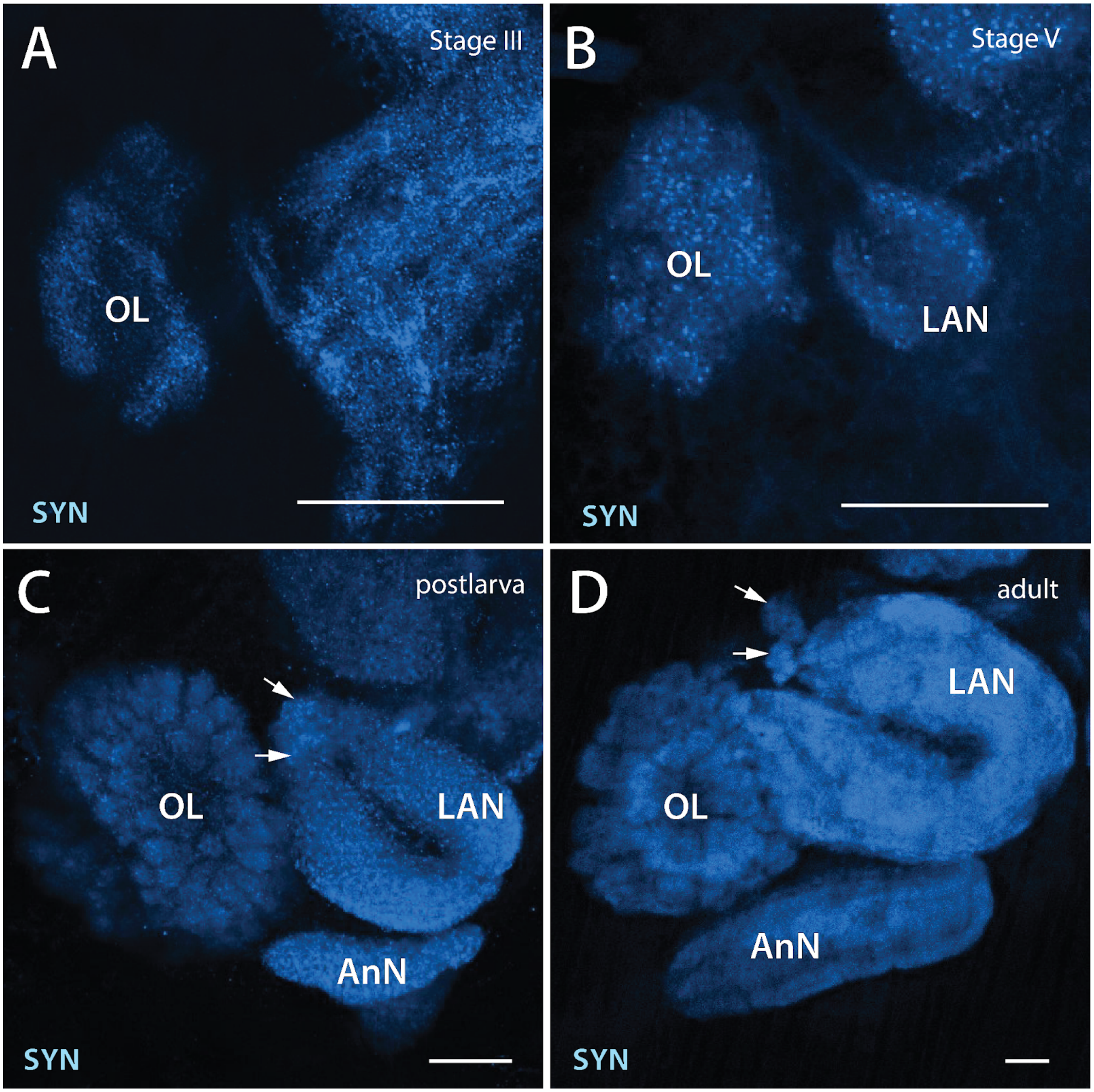


**Supplementary Figure 3.** Stomatopod sensory neuropils develop heterochronically. Both the optic lobes and olfactory lobes (OL) develop embryonically; the latter is shown here with anti-synapsin (blue) staining (Shiino, 1942). Conversely, the bi-lobed lateral antennular neuropils (LAN) and stratified antennal neuropils (AnN) appear post-embryonically. Neither sensory neuropils are present in Stage III *N. oerstedii* larvae (**A**). A LAN is present in Stage V, but it is noticeably smaller than the OL (**B**). By the transitional post-larval stage, seen here in *Gonodactylus sp.*, the LAN has increased in size and has formed two glomerular structures at the distal ends of the lobes (arrows; **C** and **D**). A small AnN is also apparent (**C**). AnN in postlarvae is unstratified, unlike AnN in adult stomatopods (**D**; *N. oerstedii*). Scale bars = 50 µm.

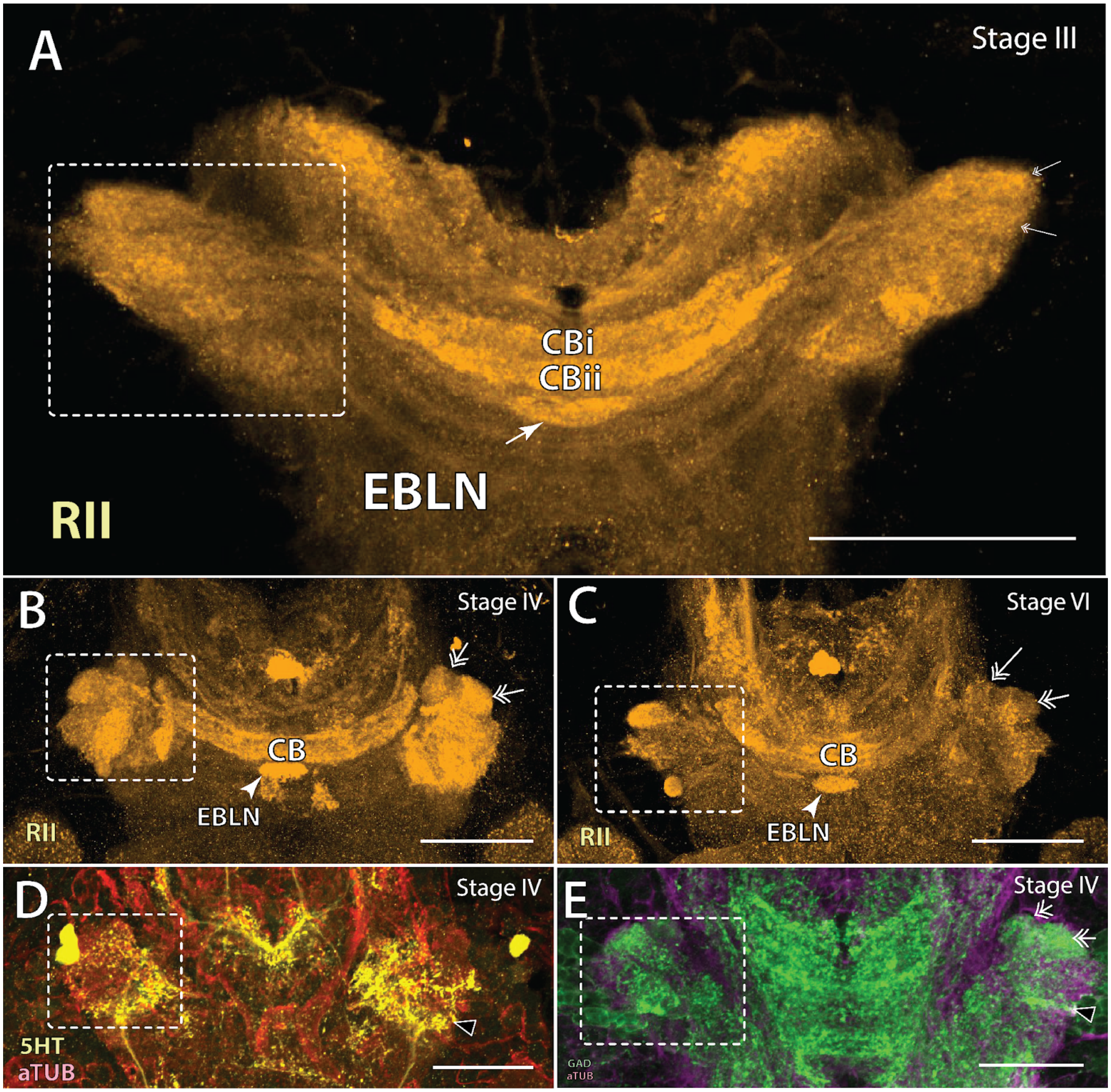


**Supplementary Figure 4.** Larval *N. oerstedii* possess multi-lobed lateral protocerebral neuropils (left structures boxed), as revealed by anti-RII (orange) immunostaining (**A-C**). These lateral protocerebral lobes (LPL) label differentially with anti-5HT (yellow; **D**) and anti-GAD (green, **E**). Equivalent lobes of this structure on the right are indicated by double arrows in **A-C** and **E**, and black triangles in **D** and **E**. Scale bars = 50 µm.
